# Pan-Cancer Drug Sensitivity Prediction from Gene Expression using Deep Learning

**DOI:** 10.1101/2024.11.15.623715

**Authors:** Beronica A. Ocasio, Jiaming Hu, Vasileios Stathias, Maria J. Martinez, Kerry L. Burnstein, Stephan C. Schürer

## Abstract

Cancer is a group of complex diseases, with tumor heterogeneity, durable drug efficacy, emerging resistance, and host toxicity presenting major challenges to the development of effective cancer therapeutics. While traditionally used methods have remained limited in their capacity to overcome these challenges in cancer drug development, efforts have been made in recent years toward applying “big data” to cancer research and precision oncology. By curating, standardizing, and integrating data from various databases, we developed deep learning architectures that use perturbation and baseline transcriptional signatures to predict efficacious small molecule compounds and genetic dependencies in cancer. A series of internal validations followed by prospective validation in prostate cancer cell lines were performed to ensure consistent performance and model applicability. We report *SensitivitySeq*, a novel bioinformatics tool for prioritizing small molecule compounds and gene dependencies *in silico* to drive the development of targeted therapies for cancer. To the best of our knowledge, this is the first supervised deep learning approach, validated *in vitro*, to predict drug sensitivity using baseline cancer cell line gene expression alongside cell line-independent perturbation-response consensus signatures.

**GRAPHICAL ABSTRACT:** 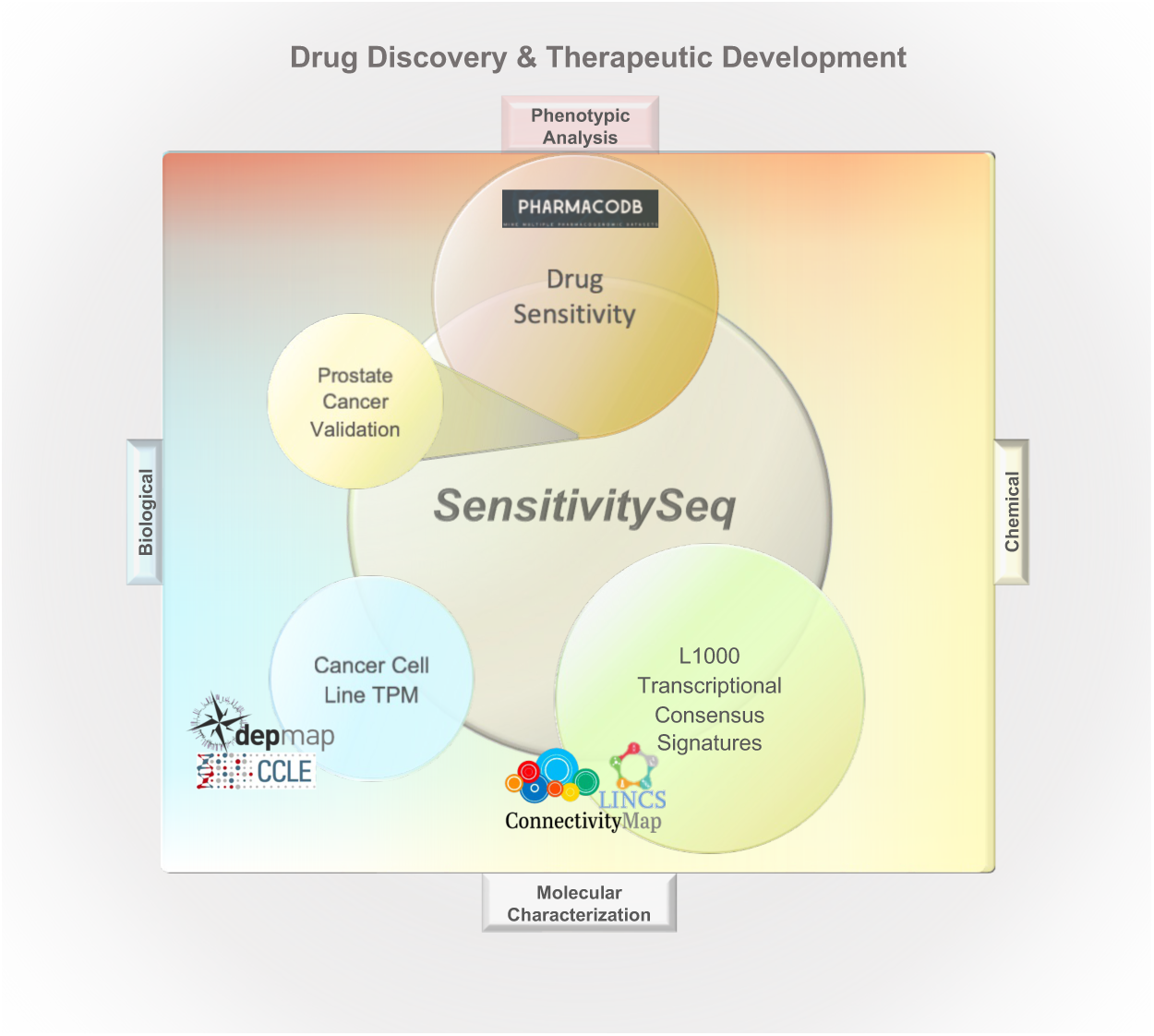

## MAIN

Cancer is a major cause of death and poses an enormous economic and societal burden worldwide^1^. As a complex, genetic disease by nature, cancer progression is unique to each patient. Additionally, tumor heterogeneity, durable drug efficacy, emerging resistance, and host toxicity remain major concerns in cancer drug development^2–4^. While there has been success in the development of targeted cancer drugs, attrition in clinical studies due to the lack of efficacy remains a major barrier. The ability to accurately predict drug response in tumor cells would enable major progress in precision oncology.

Target-based high-throughput and phenotypic cell-based screening are common approaches in the development of novel small molecule cancer therapeutics^5^. However, the de-novo development of investigational new drugs (IND) from identifying hits by high-throughput screening followed by lead optimization of biological efficacy and drug metabolism, pharmacokinetics (DMPK), and preclinical toxicity studies are frequently limited by the high upfront investment of time and monetary resources. Despite substantial preclinical investments, most compounds fail in clinical trials due to lack of efficacy. Drug repurposing and repositioning are attractive alternatives, requiring less time to approval and significantly lower development costs. However, lower expected returns on investment combined with the need to screen hundreds of compounds in pre-clinical disease models to find an efficacious drug can stifle drug repurposing efforts. Moreover, the need for anticancer drugs that simultaneously suppress tumor proliferation while minimizing host toxicity exacerbates drug discovery and development challenges further, as most early-stage drugs do not meet both criteria^3^. Pre-clinical drug discovery efforts that rely on large-scale screening can be particularly costly, time-intensive, and inefficient, especially when relying on biochemical target-based or simple *in vitro* phenotypic drug screens^6,7^.

Meanwhile, evidence increasingly supports the use of systems and computational biology-based approaches to improve the cancer drug development pipeline^8,9^. More recently, machine learning and deep learning approaches have emerged as powerful technologies to make use of the large amounts of biological profiling data that have become available in order to drive scientific discovery. Public, open-access large-scale profiling datasets and bioinformatics tools include the Library of Integrated Network-based Cellular Signatures (LINCS)^10^ and the LINCS Data Portal (LDP), the Connectivity Map (CMap) project by the Broad Institute^11,12^, the PharmacoDB drug screening database^13^, the Cancer Cell Line Encyclopedia (CCLE)^14,15^, the Cancer Dependency Map (DepMap)^16^ project, and the Genomic Data Commons (GDC) of the National Cancer Institute (NCI), which provides access to both The Cancer Genome Atlas (TCGA) and the Clinical Proteomic Tumor Analysis Consortium (CPTAC) projects. These represent some of the numerous resources that have the potential to be used to drive discovery in the cancer therapeutics research space. However, despite the increasing availability of such cancer data resources, their integration and utilization towards the development of new cancer therapeutics remains a pressing challenge. While artificial intelligence methods have been proposed as potential solutions to extract novel insights from these databases, many studies have noted the challenges posed by the limited overlap across model systems, drug treatments, compound annotations, and other biological and chemical features needed to successfully train deep learning models to predict drug response outcomes^17^. In addition, many studies have been further limited by requiring many different input features to predict drug response, ranging from molecular high throughput sequence data to chemical structures^18,19^. The physiological changes and mechanisms underlying most disease states are reflected in the transcriptome, while cellular response to treatment-induced perturbation can also be captured by changes in gene expression, as demonstrated by the CMap project and subsequent and related work^10,12^.

While multiple gene expression profile-based deep learning drug prediction models have been developed in recent years, many of these have been limited by the availability of suitable cancer treatment datasets. In particular, the development of molecular, perturbation-based cancer drug discovery models has been limited by a lack of tumor cell diversity in existing, public databases such as LINCS.

Cancer drug sensitivity is generally driven more by intrinsic tumor molecular characteristics such as gene expression than by tissue-specific cell lineage reflected by DNA profiles. While the heterogeneity of tumors lends itself to functional genomic and precision medicine approaches in cancer drug research, gene expression has been found to be a superior predictor of tumor drug response compared to other genomic features, such as genetic mutations^19,20^. This insight is also supported by evidence that gene expression is influenced by epigenetic modifications in addition to genetic mutations in tumor cells, thereby reflecting both epigenetic and genetic changes^21^.

Recent attempts to use gene expression profiles to model cancer drug response have been limited by a lack of cancer cell line representation among available transcriptional compound perturbation datasets generated under comparable experimental conditions^17^. This not only suggests utility of cell line-independent perturbation signatures, but also supports the rationale for studying individual tumor cell lines based on their unique disease signatures, rather than grouping cell lines by tissue or primary tumor type.

In this report, we leveraged the most recent LINCS L1000 datasets and further improved their robustness by generating and using cell line-independent transcriptional consensus signatures (TCS) from small molecule compounds and CRISPR genetic knockout perturbations. The TCS were used alongside baseline cancer cell line signatures derived from bulk RNA-sequencing (RNAseq) profiles to predict drug sensitivity.

Using the foundational work of the CMap L1000 project^12^, combined with the methodology used in our previous work to generate TCS for L1000 perturbations^22^, we developed transcriptional signature-based deep learning models capable of accurately predicting outcomes of drug sensitivity across cell line models of cancer, using gene expression for fewer than 1,000 protein-coding genes. Our best models were robust and highly accurate for predicting drug sensitivity in many different cancer disease states, maximizing transferability and ease-of-use by requiring gene expression values for only 969 L1000 landmark genes as input. Combined with a user-friendly web application, *SensitivitySeq* (SSeq) provides a platform to accelerate pre-clinical drug development by enabling prioritization of small molecules and dependency targets for further investigation in cancer drug discovery and development projects.

## RESULTS

### i. *SensitivitySeq* Framework and Design

To develop a pan-cancer deep learning approach that can be applied to drug sensitivity prediction for cancer cell lines, we began with initial model development and validation leading to *SensitivitySeq1.0* (SSeq1.0) models. These were subsequently updated and further enhanced in *Gen2* to obtain *SensitivitySeq2.0* (SSeq2.0) models. In the beginning of the study, diverse datasets were curated, standardized and annotated. We trained and evaluated multiple ML model architectures and designs, and thoroughly characterized these models by cross-validation. After extensive testing and analyses, two best-performing primary drug sensitivity models were further refined using an expanded dataset. Finally, after updating data sources and completing model validation analyses, an application with a searchable user interface was developed to allow researchers to query drug sensitivity predictions from a primary SSeq multi-layer perceptron (MLP) model.

To begin our investigation, we first collected, harmonized, and integrated molecular and drug screening data from multiple independent sources. In particular, PharmacoDB drug sensitivity experiments^13,23^, LINCS CMap-L1000 transcriptional perturbation signature data^10,12^, and Cancer Cell Line Encyclopedia (CCLE) gene expression profiles acquired from the Dependency Map (DepMap) portal^14–16^ were utilized as primary sources of experimental and molecular profiling data (Figures 1A and 1B). Next, we developed, trained, and rigorously evaluated a collection of initial machine learning models to develop pan-cancer, predictive DNN models and an overall approach to predict and prioritize drug sensitivity outcomes in cell line models of 28 primary tumor types, as defined by The Cancer Genome Atlas (TCGA) program of the National Cancer Institute (NCI)^24^, from 28 tissue types of origin defined by CCLE and PharmacoDB^13,23^ (Figure 1C).

**Figure 1.**
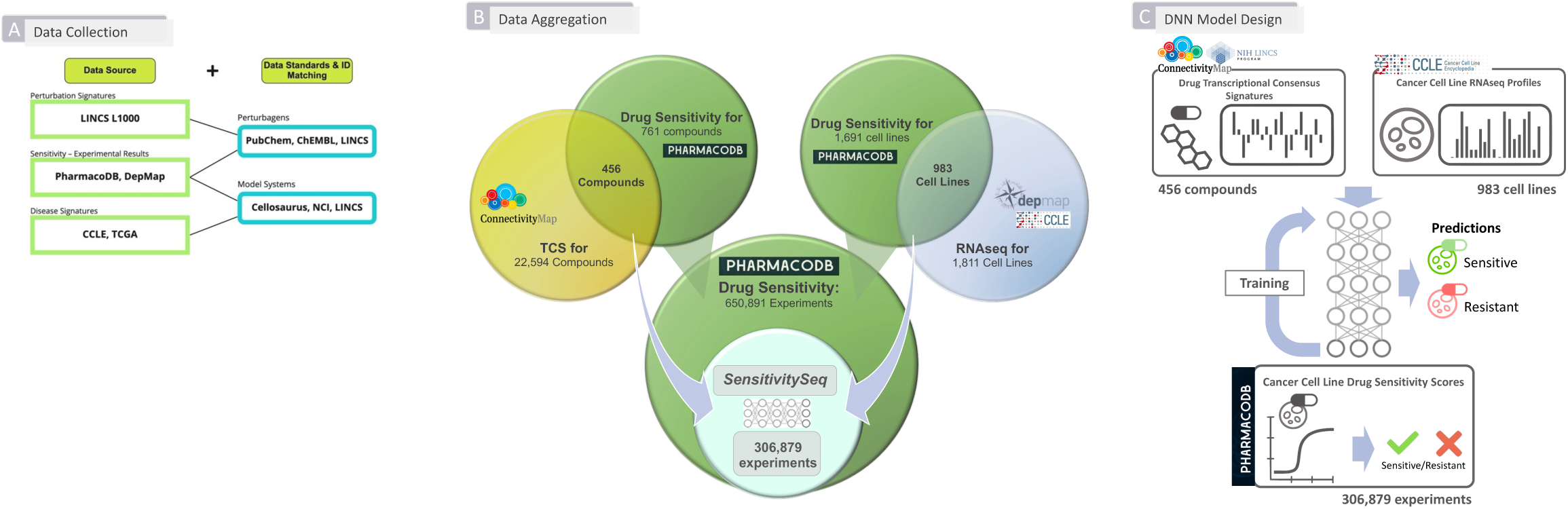
Summary of Workflow for Developing Initial *SensitivitySeq* Models. **(A)** Aggregation of data from independent sources for downstream machine learning. Three primary, independent sources of publicly available data were used to synthesize a dataset for the development of a supervised machine learning model: LINCS CMap-L1000 compound perturbation signatures, CCLE cancer cell line gene expression profiles, and PharmacoDB *in vitro* monotherapy drug response experiments. Identifiers and entities were mapped across sources using publicly available databases and metadata entries such as LINCS LDP, PubChem, and Cellosaurus. **(B) Summary of data aggregation across independent sources.** L1000 small molecules with available transcriptional consensus signatures (TCS) and CCLE cell lines were mapped to PharmacoDB drugs and cell lines, respectively, yielding 306,879 intersecting drug response experiments used to train, validate, and test the *SensitivitySeq1.0* drug sensitivity models. **(C) Overview of DNN model design and development process.** For each PharmacoDB drug sensitivity experiment with a drug sensitivity score (DSS) label, signatures for one drug and one cancer cell line model were input into the model to generate and evaluate a prediction. Binary classification was used with a prediction estimate cutoff of 0.5 to categorize drug response outcomes as either sensitive (>=0.5) or resistant (<0.5).

Following our initial model evaluation and validation, we improved our DNN models using an expanded training set, leading to our *Gen2* SSeq2.0 models. To further validate our approach, our SSeq pan-cancer drug sensitivity MLP model was applied to predict drug sensitivities for a single tissue and primary tumor type using independent, in-house generated data. This prospective validation represented a real-world application of the model unaffected by the biases of training data that often lead to overestimation of model performance based on cross-validation. In-house bulk RNAseq profiles generated for eight prostate tissue-derived cell lines, which included seven cell line models of prostate cancer (PC), were used as input to predict drug sensitivity for LINCS CMap-L1000 compounds. Twenty-six drugs were selected from SSeq model predictions for further experimental testing and subsequent validation using cell viability IC50 values against the eight prostate cell lines.

To make SSeq drug sensitivity predictions available to others, we created the *SensitivitySeq* application via Shiny, which is available online at SensitivitySeq.com and via the Schurerlab SensitivitySeq Github repository (See Methods).

### ii. Data Curation, Harmonization, Integration, and Modeling

Initial curation and aggregation of *in vitro* drug sensitivity data, small molecule transcriptional perturbation-response TCS, and CCLE cell line RNAseq gene expression profiles (Figure 1A) yielded 306,879 unique experiments measuring normalized drug sensitivity scores (DSS) from cell line-based drug sensitivity experiments, which were used to develop SSeq1.0 models. This initial aggregated dataset consisted of 456 unique small molecule compounds tested on 983 unique cell line models of cancer (Figure 1B). These datasets were further processed to harmonize and standardize the data by normalizing and scaling each dataset individually, prior to filtering steps (See Methods). Minor improvements were seen in model performance following scaling (Table S1).

L1000 compound TCS for L1000 landmark genes were chosen as the primary compound input features. Of the 978 landmark genes originally present in the L1000 data, 969 were found to overlap with the CCLE RNAseq TPM profiles. Previously, we demonstrated that compound TCS cluster by mechanism of action (MoA)^22^. A t-distributed stochastic neighbor embedding (t-SNE) analysis was performed to ensure adequate retention and representation of cancer cell line characteristics by the L1000 landmark genes, including the type of cancer each cell line models, which is associated with the CCLE Tissue Type^14,15^. This analysis showed cell lines clustered primarily by tissue type (Figure 2). In addition, a Full CCLE Transcriptome MLP model containing the 19,182 genes present in the CCLE RNAseq dataset trained, and its performance compared against the drug sensitivity MLP model based on the 969 overlapping genes. The full CCLE transcriptome input did not improve prediction performance compared to the reduced 969-gene subset (Table S1). Therefore, we proceeded with our analyses and remaining models using only the subset of the 969 landmark genes contained in both the L1000 and CCLE gene expression input datasets. These findings all support the conclusion that the core biological baseline and perturbation characteristics are retained in the reduced cell line TPM profiles and compound TCS, and together support their suitability for use as input features in machine learning models.

**Figure 2.**
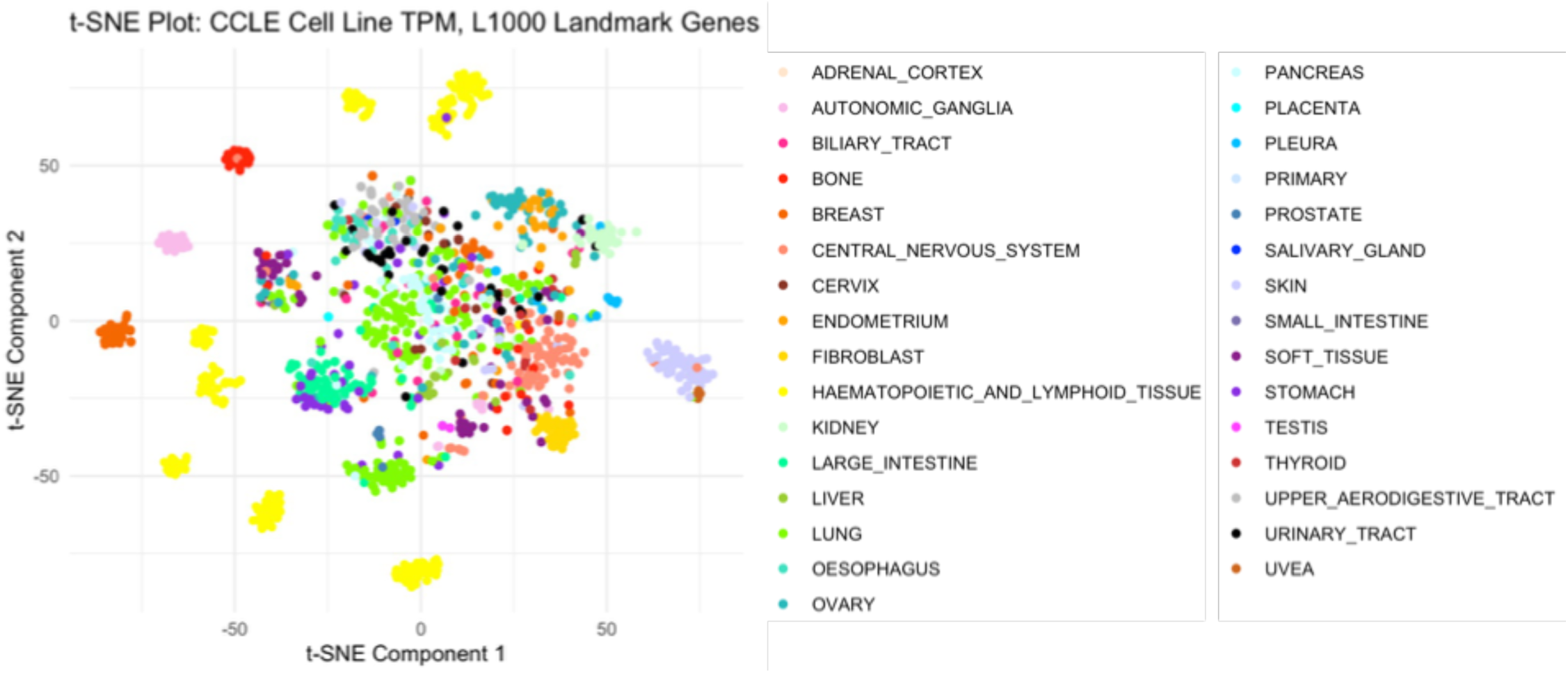
T-distributed stochastic neighbor embedding analysis of CCLE cell line gene expression signatures. CCLE Cell Line RNA-seq profiles filtered for L1000 Landmark Genes reflect cancer tissue site of origin; A t-distributed stochastic neighbor embedding (t-SNE) plot of CCLE Landmark-L1000 gene expression signatures, generated from CCLE baseline RNAseq profiles, shows cell line clustering reflects the 31 cell line tissue types of origin included in our analyses. CCLE TPM profiles were filtered for 969 overlapping L1000 Landmark genes and rescaled to a single unit range to generate CCLE gene expression signatures.

Next, a range of cutoffs across different measures of drug response in cell lines, including area above the dose-viability curve (AAC), half-maximal inhibitory concentration (IC50), and drug sensitivity scores (DSS1, DSS2, and DSS3), which represent normalized measures of the area under the dose-inhibition curve (AUC), were considered for use as a scoring metric and corresponding cutoff value. Since the aggregated PharmacoDB data consisted of data spanning multiple experimental datasets derived from independent sources, the Drug Sensitivity Score-3 (DSS3) was initially preferred as a normalized, standardized measure of drug response that reflects measures of both potency and efficacy (See Methods, Equations 1-4). Ultimately, the decision to implement binary drug sensitivity outcomes based on DSS3 values was based on the need to (1) maximize the number of experimental datapoints available to train and validate a DNN model, while (2) also providing sufficient biologically meaningful experiments with outcomes categorized as ‘sensitive’ (positive classes) in the training set to optimally train a model by allowing it to learn to differentiate between sensitivity and resistance; and (3) develop a model with retained predictive performance when a different measure of *in vitro* drug sensitivity, such as IC50, is used than the metric trained against to determine drug response outcomes. In addition, IC50 values were not available for all compounds in the datasets used due to the nature of IC50 scoring, while AAC and AUC values have also been found to outperform IC50 values for drug response prediction^25^.

Initially, a cutoff to determine binary classification of drug sensitivity was chosen based on the distributions of the DSS3 values and corresponding IC50 values (where available) that could be reasonably interpreted as sensitive, consistent with values greater than about 0.04 for DSS3, which corresponded to values less than about 5.0 micromolar (μM) for IC50 (κ = 0.5390, Z = 138.92) (See Methods, Equations 5-7). The resulting 15% of the highest scoring drug response experiments in our aggregated dataset designated as ‘sensitive’ for binary classification avoided an overly imbalanced training set and classification bias. Choosing a higher cutoff led to decreased model performance, likely due to technical bias from a highly imbalanced training set. On the other hand, choosing a lower cutoff led to worse performance, likely due to dilution of biologically meaningful labels. The robustness of this choice was later solidified further by our prostate cancer validation results based on in-house generated data (Table S11). Thus, PharmacoDB DSS3 scores reliably produced well-performing models and were found to be consistent with other measures of efficacy or potency, such as IC50, to label drug sensitivity.

Later in the study, a larger, more recent release of PharmacoDB data from PharmacoDB2.0 was integrated, which did not contain DSS3 values nor sufficient data points to calculate the DSS3 for all overlapping drug-cell line experiments to CCLE and LINCS CMap-L1000^23^. In order to maximize the size of the training dataset, PharmacoDB recomputed-AAC values, which have been normalized across datasets to adjust for variability in dose-response experiments across sources^23^, were used in place of DSS3 values, leading to our *Gen2* drug sensitivity (DS) models. For the *Gen2* DS models, a recomputed-AAC value of 27.5% was selected as a cutoff based on distribution of recomputed-AAC and corresponding DSS3 values (where available). A recomputed-AAC cutoff of approximately 27.5% corresponded to the previously used 0.04 DSS3 cutoff, with high correlation agreement seen across overlapping experiments (κ = 0.8400, Z = 724.14) (See Methods, Equations 5-7). A set of predictive performance metrics encompassing overall accuracy, precision, recall, specificity, area under the receiver operating characteristic (ROC) curve (AUROC), and the area under the precision-recall curve (AUPR) were chosen as primary evaluation criteria.

### iii. Dual-Input Deep Neural Network Fusion Model Architecture Leads to Optimal Performance

To develop the initial SSeq1.0 models, a harmonized, aggregated drug sensitivity dataset was used to train, validate and evaluate a pan-cancer DNN model capable of predicting drug sensitivity from gene expression signatures of small molecules and tumor-derived cell lines using a supervised deep learning approach. After filtering for DSS3 scores for compounds included in both LINCS CMap-L1000 and PharmacoDB that were tested on cell lines included in both CCLE and PharmacoDB, the resulting 306,879 drug sensitivity experiments were randomly divided into three separate sets. Sixty percent of the aggregated data was allocated for training, 10% of the data was allocated for post-training validation steps, and the remaining 30% of the data was set aside for downstream testing and evaluation of the fully trained model.

A series of model architectures with various hyperparameters were tested to select the highest performing model with optimal hyperparameters (See Methods). SSeq model architectures were initially designed as two identical DNN subnetworks for each input feature, followed by a merge layer and a series of dense layers (Figure 3A). A MLP architecture was constructed using two identical subnetworks, each with 969-unit input layers, followed by a series of dense layers with Rectified Linear Units (ReLU) activation. The output layer of each of the two parallel dense-layer subnetworks was then merged via a concatenation layer, followed by another series of dense layers with ReLU activation, before ending with a single node layer with a sigmoidal activation function (Figure 3B). Using the construction shown in Figure 3 as the basis of our model architecture, we arrived at the final architecture after a series of modeling tests and evaluations to assess the impact of numbers of layers and nodes, as well as activation functions, optimizers, number of epochs, and other model specifications (See Methods).

**Figure 3.**
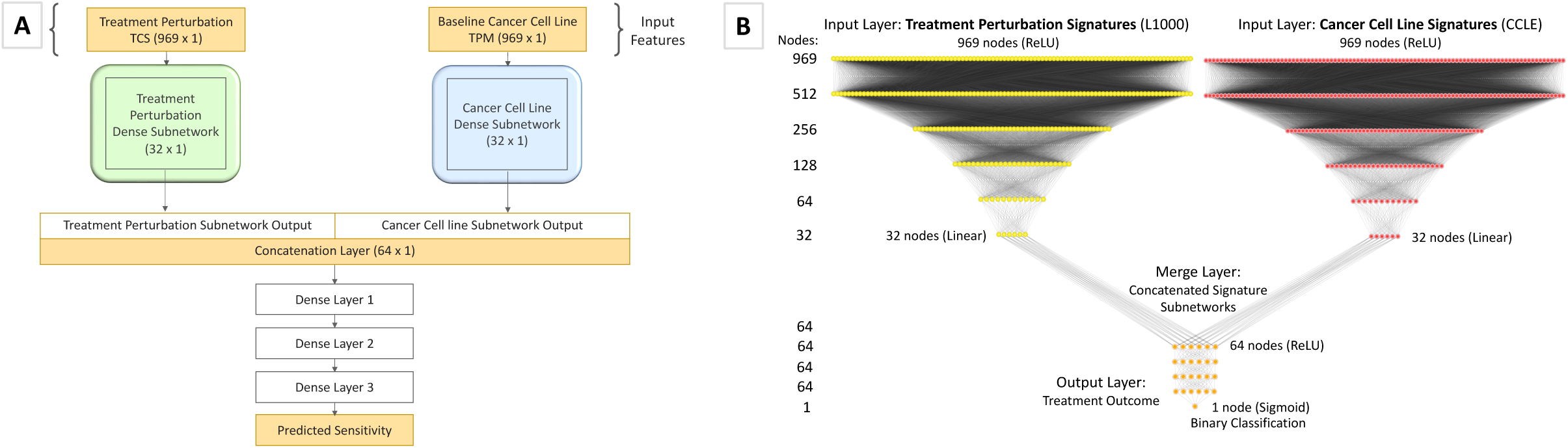
SSeq DNN Architecture and Design. **(A)** Abstract description of the main model architecture and input data used to develop SSeq models. The model architecture is composed of initial dual feature subnetworks with a series of dense, hidden layers, followed by a merge layer, and another set of dense, hidden layers after concatenation. The final layer binary output specifies whether a specific treatment perturbation (e.g. induced by a small molecule) is predicted to result in inhibited proliferation of a given cell line. **(B)** The detailed architecture used for the SSeq MLP Drug Sensitivity and Genetic Dependency models is shown. Dual subnetworks are composed of 969 nodes in the input layer of each subnetwork with a ReLU activation function, followed by a series of four hidden layers with ReLU activation functions and 512, 256, 128, and 64 nodes, in descending order. The last dense layer of each subnetwork features 32 nodes and linear activation. The output of each subnetwork is subsequently concatenated via a merge layer with 64 nodes and ReLU activation. After three more 64-node dense, hidden layers, the transformed output is passed to a final layer composed of a single node with a sigmoid activation function for binary categorization.

Binary encoding and classification were employed using PharmacoDB normalized, AUC-based drug sensitivity DSS3 scores with a cutoff of 0.04 to define drug sensitivity of cell lines to LINCS CMap-L1000 small molecule compounds based on the respective gene expression signatures of each. After training the binary classification model, the left-out test set was used to evaluate the performance and accuracy of the prediction estimates of the model. A binary classification model was chosen to facilitate broader model applicability, whereby prediction estimates are generated based on the probability of drug sensitivity as defined by the sensitivity cutoff, rather than quantitative prediction estimates based on the output label DSS3 value, which would require further conversions to translate predictions to other metrics and reduce the amount of usable data.

Binary scores determined by prediction estimates were used to evaluate the performance of the trained DNN models. A prediction estimate cutoff of 0.5, indicating a 50% probability of meeting criteria for ‘sensitive’ (i.e. a score of ‘1’) was used to determine sensitivity categorization for positive classes. Prediction estimates greater than or equal to 0.5 were categorized as a ‘1’ (sensitive), and prediction estimates less than 0.5 were categorized as ‘0’ (resistant). Using these cutoffs to evaluate model performance in predicting outcomes for the test set, different model architectures were tested spanning multi-layer perceptron (MLP) models, one-and two-dimensional convolutional neural network (CNN) models, and various combinations of these, before settling on two highest-performing model architectures. Our initial primary pan-cancer MLP drug sensitivity model displayed 90.5% accuracy, 93.1% AUROC, 95.7% specificity, 62.8% recall, and 73.5% precision (Table S1). In addition, we retained a secondary model with a two-dimensional CNN (2D-CNN) (Figure 5). The 2D-CNN model showed slightly lower overall accuracy (89.1%) and AUROC (91.2%), with a higher recall (71.6%) at the apparent cost of reduced precision (63.9%) and slightly reduced specificity (92.4%) (Table S2). The 2D-CNN model architecture slightly outperformed the 1D-CNN (Table S2). Later, class weight optimization was performed on each of the two retained models using a 5:1 ratio of positive to negative classes. While performance remained similar for the MLP model, the 2D-CNN with class-weight optimization (CW) showed increased recall and AUPR (Table S2) and was thus retained in place of the earlier 2D-CNN (without class weight optimization) as a primary SSeq model.

### iv. Cross-Domain Validation Reveals *SensitivitySeq* Models are Robust

In order to ensure model robustness and applicability, extensive cross-domain validation and evaluative analyses of the DNN models and model architectures were completed. Multiple validation studies were performed, including 10-fold cross-validation, very low training to test ratios, and evaluation of compound and cell line cross-domain applicability. Monte Carlo repeated randomized sub-sampling and 10-fold cross-validation analyses showed that our modeling strategy was robust and retained performance even when training set sample sizes were small (Tables S3, S4, and S5). The results of these analyses also suggested that increasing the available data set size would likely improve model performance, which was later confirmed by our *Gen2* models (Table 1).

**Table 1.** Comparison of Performance for Initial vs *Gen2* Finalized Deep Neural Network Models. Model predictions were evaluated based on several measures of model performance. The first model listed represents the performance for our initial, SSeq1.0 pan-cancer drug sensitivity MLP model, trained and evaluated using scaled input datasets. Results for the updated versions of the SSeq2.0 MLP and 2D-CNN models, after adding more than 150,000 additional experiments from recently released open-source datasets to the training set, are shown in row 2 and row 4, respectively. 5:1 class-weights were applied as a balancing strategy to adjust for an imbalanced training set and retained for the finalized 2D-CNNs. Increased recall was seen in the MLP model following training with the expanded *Gen2* dataset, while increased precision was seen in both the 2D-CNN model and the MLP. Higher AUPR was recorded for both updated, finalized SSeq2.0 models.

| Model | Loss | Accuracy | AUROC | AUPR | Precision | Recall | Specificity |
| --- | --- | --- | --- | --- | --- | --- | --- |
| **SSeq1.0 Drug Sensitivity MLP** | 0.2264 | 90.54% | 93.07% | 76.17% | 73.46% | 62.84% | 95.74% |
| **SSeq2.0 Drug Sensitivity MLP** | 0.1917 | 92.66% | 95.06% | 83.04% | 75.28% | 75.76% | 95.63% |
| **SSeq1.0 Drug Sensitivity 2D-CNN** | 0.2691 | 88.84% | 93.44% | 76.94% | 61.08% | 80.93% | 90.32% |
| **SSeq2.0 Drug Sensitivity 2D-CNN** | 0.1963 | 92.26% | 95.35% | 82.33% | 71.18% | 80.98% | 94.24% |

To assure classification predictions were based on the gene expression signatures, we scrambled the sensitivity labels while maintaining the ratio of classes. Subsequently, we repeated all training and validation steps used for our initial SSeq1.0 MLP model. Cross-validation AUROC values were measured to be approximately 50% for all architectures tested, indicating random classification as expected (Table S1; Figure 6). The performance of the MLP model trained with randomized labels highlights the importance of evaluation based on multiple criteria particularly in the case of imbalanced datasets; all predictions were negative, resulting in 0% precision and 0.5 ROC, but 100% specificity and decent accuracy, due to the overrepresentation of negative classes appearing as the only detectable pattern in the training set when labels were scrambled (Table S1). Overall, these results suggested that the model architectures and datasets used did not lead to overfitting.

Next, cross-domain validations were performed for cancer cell lines grouped by tissue types. Models were trained excluding one tissue followed by predicting sensitivities for the tissue group that was not included in the training data. Results showed that the models can accurately predict drug sensitivity for novel tissue types, which they had not been trained on, supporting our pan-cancer approach (Figure 4; Table S6). The cross-domain validation prediction accuracy was greater than 78% (Figure 4C) with an AUROC above 80% (Figure 4D) across all cell line tissue types despite the range in representation across the training data (Figures 4A and 4B). Similar cross-domain validation was performed for compounds based on chemical structure scaffold (chemotype) and ligand type (Extended Data Fig. 1 and Extended Data Fig. 2; Table S7). For chemotype cross-domain validation, compounds were clustered based on topological features (FCFP4 and ECFP4 fingerprints) (See Methods). Models were trained on all but one cluster, and predictions were then made for the left out chemotype (Extended Data Fig. 1). Overall, results showed that the models are capable of predicting drug sensitivity for novel chemotypes for which no data were present in training.

**Figure 4.**
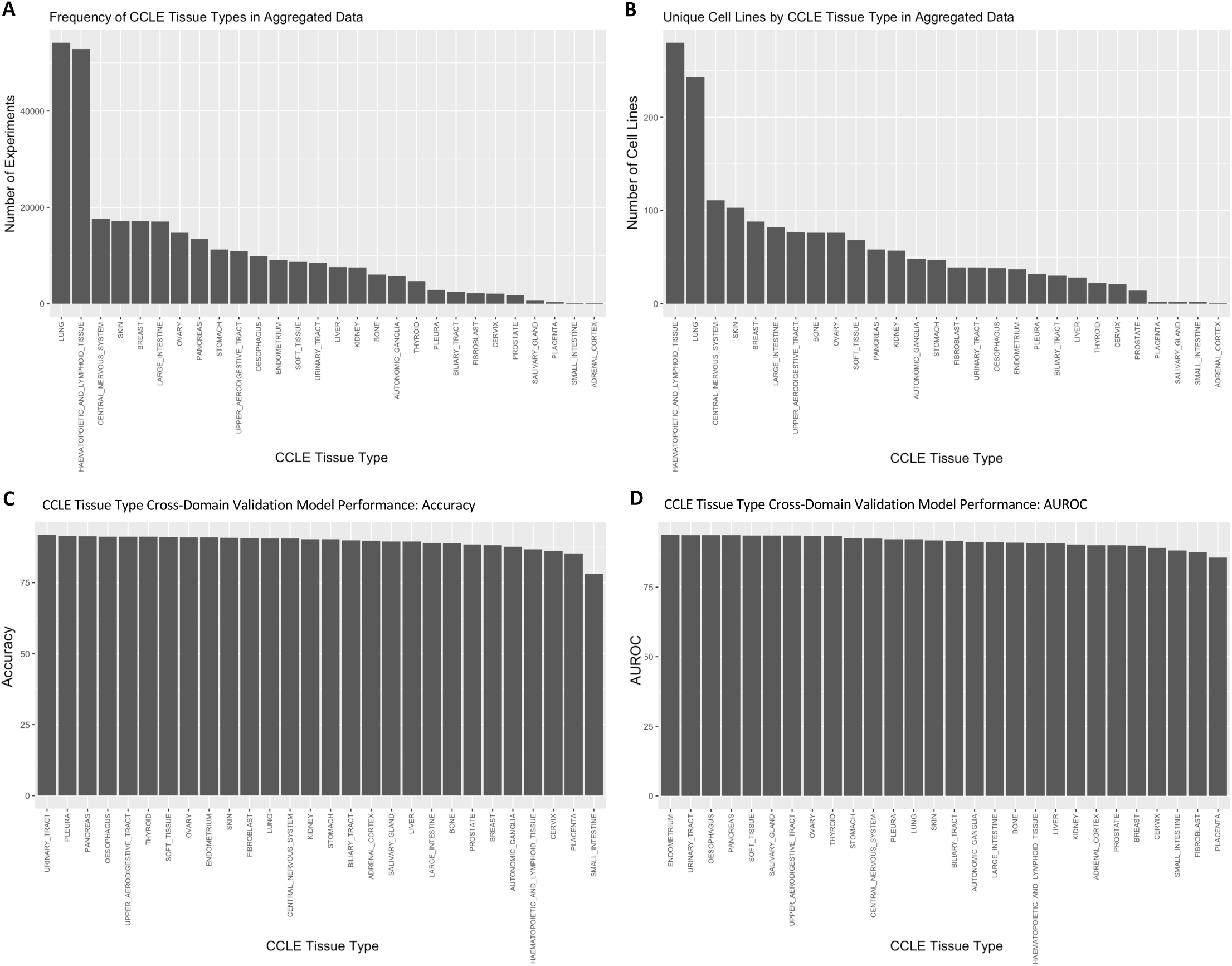
Number of drug sensitivity data points by CCLE Cell Line Tissue Types in aggregated training data vs. prediction performance by tissue type. (A) The frequency of experiments for CCLE tissue type is shown for cell lines in the initial drug sensitivity aggregated training dataset. Number of experiments refers to the number of unique drug-cell line combinations with a DSS3 score recorded and applied as a training label. (B) The frequency of cell lines corresponding to each CCLE tissue type is shown for cell lines in the initial drug sensitivity aggregated training dataset. (C) The cross-domain validation accuracy for cell lines grouped by tissue type is shown for each category. The prediction accuracy was high (>78%) across all cell line tissue types despite the range in representation across the training sets. (D) The cross-domain validation AUROC for cell lines grouped by tissue type is shown. Like accuracy, the prediction AUROC was also high (>80%) across all cell line tissue types.

To address our imbalanced training labels, in which only approximately 15% of experiments met criteria for ‘sensitive’ (positive class) categorization without excluding significant amounts of data, we performed further analyses including calculating the AUPR and testing approaches to balancing the model. Resampling of the data by splitting the overrepresented ‘resistant’ groups into five separate training sets was evaluated. These five groups of ‘resistant’ data were each combined with the ‘sensitive’ set of data to create smaller, balanced training sets and used to train individual, stand-alone models. The five balanced training sets were also used to develop a serial model, where initial training with one balanced dataset was followed by consecutive re-training with each of the remaining four balanced datasets (Table S8). However, manipulating class weights to accommodate the full, imbalanced training set led to better prediction performance than splitting training data into balanced subsets and training models with reduced amounts of data, and almost identical performance to the serial model (Table S8). The CW 2D-CNN model (Figure 5) showed better performance compared to the MLP, so this model was retained despite its higher complexity (Tables 1 and S8). The 2D-CNN model architecture also slightly outperformed the 1D-CNN (Table S2). Additionally, as AUROC can be skewed for imbalanced data, AUPR was determined for each model. The AUPR of our SSeq1.0 MLP model was 76.2%, while the AUROC was 93.1% (Table S1; Figure 6).

**Figure 5.**
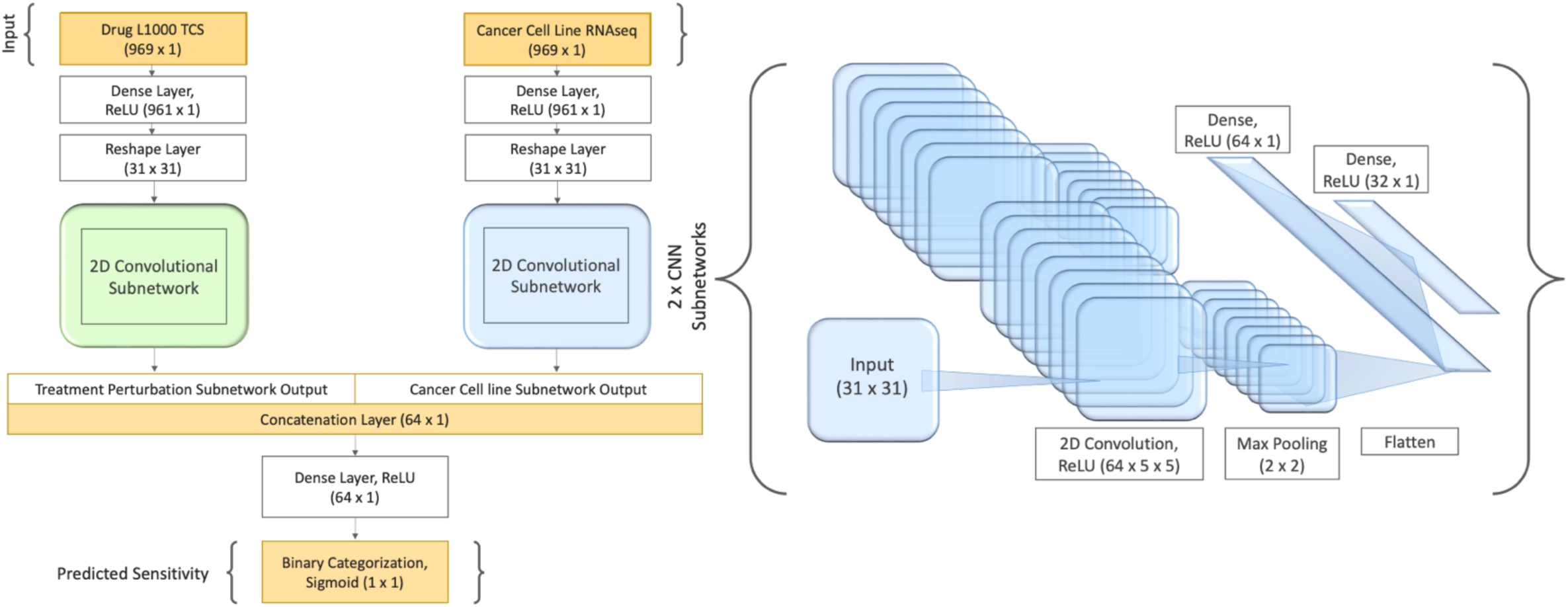
Model Architecture and construction of two-dimensional convolutional neural network model, SSeq CNN. A two-dimensional convolutional neural network (2D-CNN) model was constructed and implemented with 5:1 class weight applied as a balancing strategy and hyperparameter optimization method and was compared to our SSeq1.0 MLP drug sensitivity model. The architecture of each of the two identically constructed 2D-CNN subnetworks is shown on the right. The final version of the 2D-CNN is more complex, balanced, and displays higher recall compared to the SSeq MLP model, as shown in Table S2.

**Figure 6.**
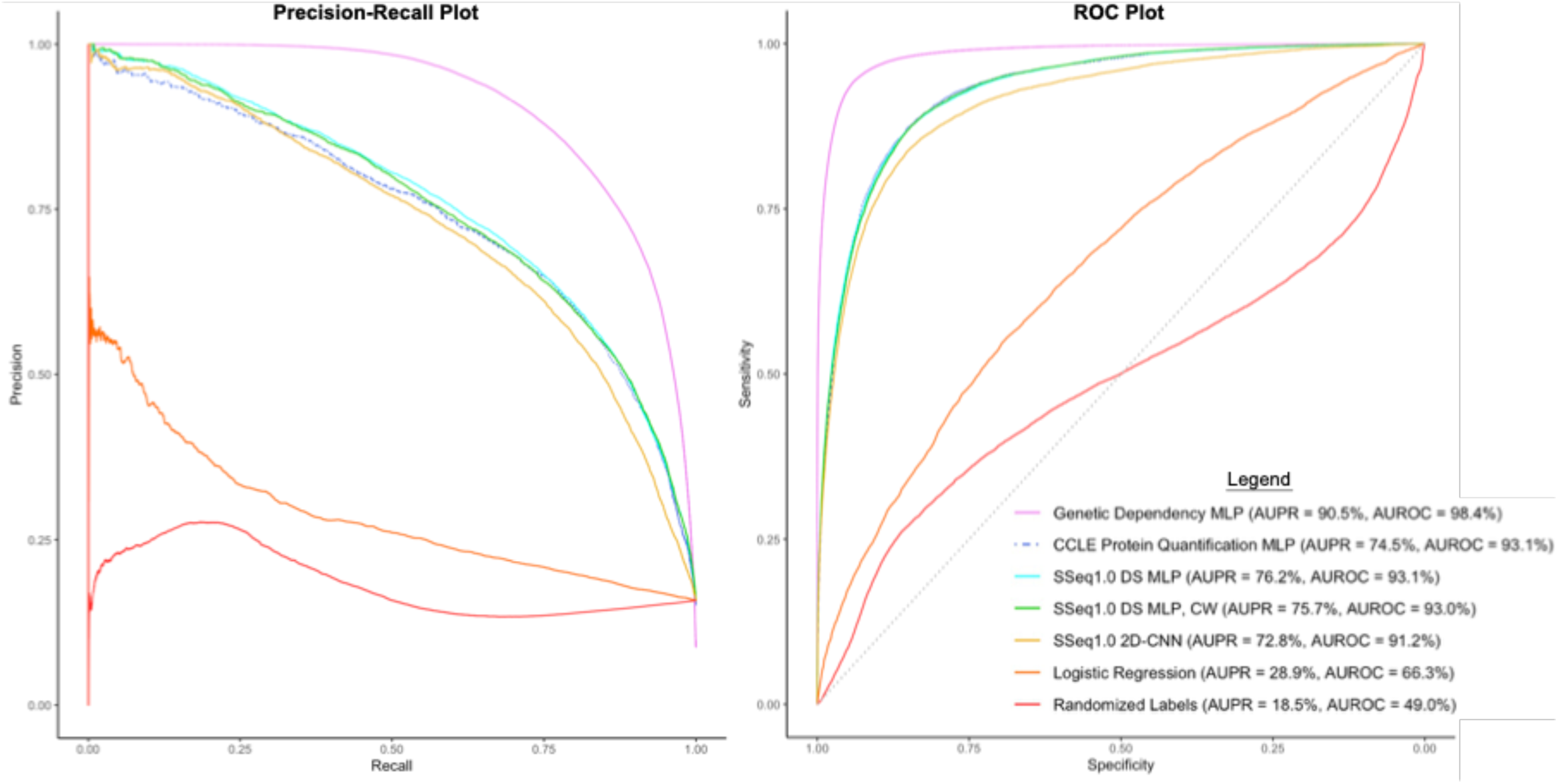
Precision-Recall curve and ROC curve plots for seven machine learning models evaluated. The legend shown applies to both plots and specifies the area under the precision-recall curve (AUPR) and the area under the ROC curve (AUROC) for each model. The Genetic Dependency MLP model displayed the highest performance according to these metrics, with an AUPR of 86.5% and an AUROC of 95.9%, indicating reliable prediction performance. The measured AUPR for the SSeq1.0 drug sensitivity (DS) MLP model was 76.2% and the measured AUROC was 93.1%, while the SSeq1.0 MLP with class weight optimization (SSeq1.0 DS MLP, CW) performed similarly with an AUPR of 75.7% and AUROC of 93.0%. All DNN models displayed higher measured AUC values than both the MLP model trained with Randomized Labels, and the basic Logistic Regression model.

A range of different machine learning algorithms and DNN architectures were implemented and tested to compare their predictive performance (Table S2). First, a basic logistic regression model was trained and tested to serve as a benchmark to evaluate the performance of our more complex DNN models (See Methods, Equation 8). The binary classification, multivariate logistic regression model did not distinguish any positive outcomes. All prediction estimates generated were below 0.5, resulting in only predicted scores of ‘0’, indicative of a treatment-resistant outcome (‘resistant’ classification). This contributed to 0% precision and undefined recall, suggesting that this algorithm simply did not work. This was likely due to the inability to normalize the input data across different measures (TPM and TCS signatures), in contrast to the DNN models. Native handling of multi-modal data is a fundamental advantage of deep neural networks, and these results support the approach of using more complex MLP and DNN models for this data. The performance results of these models are summarized in Table S2.

Single-feature models containing only one of the two primary input features used in our primary modeling approach were constructed to assess predictive performance for each feature and feature-subnetwork individually, as well as to validate our dual feature-subnetwork approach. These included DNN models trained and evaluated using only PharmacoDB drug sensitivity labels as output and, separately, (1) LINCS CMap-L1000 small molecule transcriptional consensus signatures (TCS), and (2) CCLE cancer cell line RNAseq TPM profiles as isolated input features. The results of this analysis showed that LINCS CMap-L1000 small molecule TCS were reasonably accurate predictors of drug sensitivity across all cell lines. CCLE profiles alone did not perform highly, likely due to the contradicting results of sensitive and resistant outcomes for different cell lines without corresponding treatment perturbation features from TCS. L1000 TCS were quite strong predictive features of *in vitro* drug sensitivity on their own. However, including both L1000 TCS and CCLE signatures as input features led to higher performance metrics, including AUROC and recall for both MLP and CNN models (Table S9). Thus, small molecule TCS contribute most of the information to predict drug sensitivity, but are not sufficient by themselves; in particular, CCLE cancer cell line gene expression signatures would be required as input features to predict cell line-specific drug sensitivity.

Our primary interest was to develop highly predictive models of small molecule-induced drug sensitivity. We therefore introduced small molecule topological features (specifically, extended connectivity fingerprints of length 4, ECFP4) as an additional modality to test if our SSeq MLP model would achieve enhanced performance through the incorporation of additional chemical structure information. ECFP4 are circular topological fingerprints that encode structural information based on atom neighborhoods within a radius of 4 around each atom in the molecule and are commonly used as chemical structure topological descriptors, including in our recent study^26^. Upon assessing model performance with ECFP4 fingerprints added as a third input feature subnetwork to our drug sensitivity SSeq MLP model architecture, with a modified first layer to accommodate a 1024-unit input vector, we found that the addition of ECFP4 did not lead to improved model performance compared to our dual-feature drug sensitivity SSeq MLP model with L1000 small molecule TCS and CCLE input features. Therefore, the level of performance enhancement seen with the inclusion of ECFP4 fingerprints as a third feature network in our drug sensitivity model did not justify the increased complexity.

We also conducted a separate assessment in which an adaptation to our SSeq MLP model was made by substituting an ECFP4 fingerprint feature subnetwork for the L1000 TCS, leading to a new ECFP4-CCLE MLP model. The model containing ECFP4 fingerprints in place of compound TCS did not outperform the drug sensitivity MLP model developed with L1000 TCS and CCLE input features (Table S10). Altogether, the results of our ECFP4 analyses suggest that ECFP4 descriptors provide some level of predictive capability, while combining them with compound TCS does not appear to yield significant enhancements.

### v. *SensitivitySeq* Model Architecture Shows Extended Applicability

After validating our drug sensitivity models and prediction pipeline, we tested the applicability and utility of our methods to other datasets relevant to cancer drug discovery. First, we trained a new model by adapting the architecture used to develop our SSeq DS MLP model with CCLE protein quantification data^27^ in place of RNAseq data as cancer cell line signature input. Our protein quantification MLP model displayed slightly higher recall than our SSeq1.0 MLP with CCLE RNAseq signatures, but also showed slightly lower precision (Table S9). The results of using protein quantification features as model input imply that the DNN-based machine learning models and architecture generalize to accommodate molecular feature data beyond messenger RNA (mRNA) transcription-based signatures.

Next, we developed a genetic dependency model applying the same model architecture and comparable methods to our drug sensitivity model, using L1000 CRISPR perturbation TCS and Achilles genetic dependency data, alongside CCLE RNAseq signatures, similar to our drug sensitivity models. A gene dependency score cutoff of 0.5 or greater was applied to designate positive classes corresponding to a probability greater than 50% of a given cell line exhibiting dependency on a given gene. The input dataset underwent a randomized split of 60-10-30, allocating 60% for training, 10% for validation, and 30% for testing to evaluate the finalized model. Our genetic dependency model showed impressive performance across nearly all metrics, except for precision (Table S9). The results of the genetic dependency model helped to cement the conclusion that our methodology and modeling techniques are robust and applicable to forms of input features and perturbation-response TCS beyond those elicited by small molecule compounds.

### vi. *SensitivitySeq* Performance Improved with an Expanded Training Set

Finally, we updated our training datasets for our drug sensitivity models with the latest releases of each of the three main datasets used to develop the initial models (LINCS CMap L1000, CCLE, PharmacoDB datasets) leading us to the development of our *Gen2* models (see Methods). The large datasets used to train the *Gen2* models contained over 485,000 and 4.5 million unique experiments for the drug sensitivity and genetic dependency models, respectively (Extended Data Fig. 3). The improved performance results of our drug sensitivity models following re-training with the expanded datasets are summarized in Table 1. Combined with the Monte Carlo and *k*-fold cross-validation analyses undertaken on our initial SSeq models (Tables S3, S4, and S5) these results support the conclusion that larger training set sample sizes lead to higher performance across multiple metrics. Therefore, continuing to expand our models as more data becomes available will likely lead to more reliable model performance in the future.

### vii. *SensitivitySeq*-Predicted Compounds Validated in Prostate Cancer

To further study the applicability and utility of our models, we applied the pan-cancer DNN SSeq2.0 MLP model to a separate, independent set of in-house generated prostate cancer experimental data, consisting of (1) prostate cell line RNAseq profiles and (2) *in vitro* drug response screens (see Methods). We generated baseline RNAseq profiles using normalized transcripts per million (TPM) data for eight human cell lines derived from prostate tissue, in conjunction with drug sensitivity screens performed on the same eight cell lines. These included seven prostate cancer (PC) cell line models: *22Rv1*, *C4-2B, DU145*, *LNCaP*, *NCI-H660, PC3*, *VCaP*, and one non-tumorigenic prostate cell line: *RWPE-1*. Applying the SSeq2.0 MLP model, drug sensitivity predictions were generated for the eight prostate cell lines to 7,398 small molecules from the LINCS CMap-L1000 perturbation TCS data. From this set, 208 unique compounds were predicted by the model to be active against one or more PC cell lines, and 26 compounds from this list were selected to test *in vitro* in prostate cell lines. An IC50 cut-off concentration of 5.0 µM was implemented to determine cell line sensitivity to each of the 26 compounds.

Evaluating SSeq2.0 model predictions for prostate cell lines against the binarized in-house experiment IC50 values (using a cut-off of IC50 < 5.0 µM indicating sensitivity) revealed 23 out of 26 (88.5%) compounds correctly identified as active against preclinical cell line models of prostate cancer. The performance and evaluation results for this prospective, independent prostate cancer evaluation of our SSeq MLP model are summarized in Tables S11 and S12.

*In vitro* drug response tests showed positive drug sensitivity results in at least one prostate cancer cell line for 23 of 26 drugs selected from the list of positive SSeq2.0 model predictions, as defined by an IC50 concentration less than 5.0 µM (Table S12). SSeq1.0 was evaluated for comparison purposes, with results showing more cell lines for each compound, on average, exhibited experimental sensitivity than predicted by the model. Thus, the SSeq1.0 model underperformed for recall with a substantial number of false negative predictions, while this was improved in SSeq2.0. In both models, the majority of positive predictions made by each model were found to be true positives, with drug sensitivity exhibited *in vitro* (Table S11).

### viii. *SensitivitySeq* Application

To make the model predictions readily available to others, we developed a user-friendly Shiny-based application*. SensitivitySeq* provides a clean user-interface to access and query a virtual library of *in vitro* monotherapy drug sensitivity predictions for all CCLE cell lines treated with each LINCS CMap-L1000 compound. *SensitivitySeq* allows users to query sensitivity predictions for a CCLE cancer cell line model treated with a LINCS L1000 compound of interest from our pan-cancer DNN model prediction estimates. As of the publication date, the first version of *SensitivitySeq* is currently available via https://sensitivityseq.com and via the *SensitivitySeq* GitHub repository (https://github.com/schurerlab/sensitivityseq). All future updates and expansions to *SensitivitySeq* will be documented on the GitHub page.

## DISCUSSION

We have developed a transcriptional signature-based, pan-cancer DNN model capable of predicting *in vitro* drug response outcomes for individual tissue-types and cell line models of cancer using gene expression quantification for fewer than 1,000 genes as input. Our vigorous analyses and model evaluations allowed us to develop an efficient, user-friendly model requiring minimal input, processing time, and computing power, which performs as well as more complex models.

Although prostate cell lines were underrepresented in our training data, as shown in Figures 4A and 4B, the pan-cancer drug sensitivity MLP model performed reasonably well when PC cell line data from a separate source was used as input. Although the cell line-specific prediction accuracy for the 26 compounds selected from positive model predictions was lacking, 23 of 26 compounds were found to inhibit growth in at least one tumorigenic PC cell line with an IC50 less than 5.0 µM. The remaining three compounds each showed an IC50 between 5.0 µM and 15.0 µM in at least one tumorigenic PC cell line. While some PC cell lines were incorrectly predicted to be sensitive or resistant to select compounds and were experimentally disproved, the model performed very well overall for prioritizing effective compounds for prostate cancer, especially considering the low representation in the training set (Figures 4A and 4B). Therefore, this pan-cancer model was shown to be effectively implemented as a tool with which to pre-screen therapeutic candidates *in silico* for individual cancer indications, prior to further laboratory testing and development.

We used a tissue-based validation analysis to show that our method is applicable for predicting drug response in cell line models of tissue types not included in the training data (Figure 4; Table S6). In addition, we also performed compound-based validation analyses to show that our methods are not biased toward any specific classes of chemical compounds (Extended Data Fig. 1 and Extended Data Fig. 2).

Finally, we created multiple adaptations and variations of our initial drug sensitivity MLP model, resulting in new models trained on multiple types of biological and chemical input features to predict both drug sensitivity and genetic dependency. Collectively, the sustained model performance metrics seen across the multiple features and model variations tested support the conclusion that our methodology is robust and applicable to many different datasets. The application of our modeling methods to cell line protein quantification input features and to ECFP4 fingerprint input features, in addition to LINCS CMap-L1000 CRISPR perturbation TCS to predict gene dependencies, altogether show that our methodology has high transferability and applicability to predicting perturbation response outcomes in cancer cell lines beyond transcription-based small molecule drug response.

LINCS CMap-L1000 transcriptional signature data for small molecule compound perturbagens were used to generate TCS that would function as dose-and cell line-independent representations of biological response to perturbation. Although the subset of LINCS CMap-L1000 compound perturbation data we used contained data for 230 cell lines, only a small subset of these cell lines was treated with each compound perturbagen. Therefore, not every compound was tested on the same cell lines. In addition, a range of treatment concentration doses were used to treat cell lines to generate measured perturbation responses. These doses ranged from 4.115×10^-5^ to 3.162×10^5^ µM in the subset of data we used.

Our drug sensitivity models rely on cell viability and drug response assay data with drug sensitivity scoring metrics derived from dose-response curves and used to determine binary training and test set labels. These data originate from multiple sources, presenting the challenge of determining a standardized measure of drug sensitivity. The DSS3 and recomputed-AAC are both derived from a normalized transformation of the dose-inhibition AUC^13,28^ and were chosen as the drug sensitivity metrics to determine the binary outcome labels used to develop the initial and *Gen2* PharmacoDB drug sensitivity models, respectively, in part to accommodate the multi-source training dataset.

Additionally, an imbalance of positive and negative classes was another limitation of the training data we used. To address this, we evaluated several strategies including resampling and adjusting class weights. Overall, most of the class-balanced models showed higher recall, but generally reduced measures of precision, specificity, and overall accuracy (Table S8). Generally, small molecule bioactivity data is expected to be imbalanced because most compounds are not active against most targets, and the models should incorporate these characteristics.

Our SSeq1.0 MLP model was highly specific, but not as sensitive as other models (Tables 1, S1, and S2). While we created other models that reflected higher sensitivity upon evaluation, the improved recall seemed to come at a cost of lowered precision (Tables 1, S1, and S2). Overall, our *SSeq* MLP drug sensitivity model is better suited for eliminating compounds that are unlikely to inhibit cell growth for a cell line model of interest and is best suited for reducing the number of compounds *in silico* that would need to be tested via downstream *in vitro* experiments. Thus, the model is highly accurate in identifying ineffective compounds for a particular cell line as true negatives; Less than 5% of screened sensitivity assays (i.e. a single cell line treated with a single compound) are expected to be wrongly categorized as sensitive when actually resistant. However, while the false positive rate is low, as a trade-off, the model is also more likely to produce false negatives. In addition, the actual rates for false positives and false negatives vary by cell line, and in turn, by tissue of origin and cancer type, which may be due in part to sequencing read quality and is also likely influenced by amount of representation in the training dataset (Figure 4).

Despite these limitations, we expect our predictive pan-cancer DNN models and *SensitivitySeq* to be useful for *in silico* screening to reduce the number of compounds requiring *in vitro* tests. Since our model has been developed using a limited number of features, relying entirely on 969-gene subset of the human transcriptome, we expect it to be especially useful as a tool used by researchers in conjunction with other methods to select a limited number of compounds for further testing.

In addition, ECFP4 fingerprints were generated using SMILES sourced from LINCS metadata for overlapping L1000 and PharmacoDB compounds common between both datasets. At the time of this publication, only 421 of these shared compounds were represented by SMILES in the LINCS database, and the resulting difference in training set size may have led to lower performance than would be expected with a more complete training set. Indeed, model performance for our drug sensitivity MLP with L1000 TCS and CCLE TPM input features using a comparable training set containing only 421 compounds showed comparable measures of performance to the ECFP4-CCLE model (Table S10). While adding chemical structure features using ECFP4 fingerprints in a third subnetwork alongside L1000 TCS and CCLE RNAseq signatures did not appear to improve drug sensitivity prediction performance enough to justify the added complexity, the creation of a separate model developed to predict drug sensitivity from ECFP4 fingerprints and CCLE RNAseq data is arguably well justified by the highly likely scenario in which a researcher has only compound SMILES available rather than L1000 profiles and wishes to pre-screen compounds against cell lines *in silico*. While beyond the scope of this current study, we plan to explore adding an ECFP4-CCLE drug sensitivity model to our *SensitivitySeq* suite in future deployments.

There are many potential ways our models described in this study could be improved, and we plan to investigate these possible avenues of improvement in future studies. For example, one limitation for this study was the amount of available data after filtering for overlapping data across three individual sources. As more data are added to these and other databases, we expect greater amounts of data synthesis to be possible, as well as improved training outcomes with the availability of larger, improved datasets. In addition, adding more features may be another way to improve model performance. However, adding more features often requires more data to be filtered out when matching entities and identifiers across sources, as seen in our ECFP4 model analyses. In the future, as more data becomes available and curation efforts continue, we plan to investigate within this area further. Our drug sensitivity prediction models were extensively validated for use with small molecule drugs given the large amount of curated data available for these compounds. However, our limited analyses on adding chemical and structural information as features showed that the incorporation of these did not significantly improve model performance. It is likely also reasonable to expect our modeling approach to be applicable to predicting cell sensitivity to cancer treatments beyond small molecule drugs, so long as these induce a measurable transcriptional perturbation response. Although our drug sensitivity models were developed using a pan-cancer approach with small molecule perturbations and cell line models, we expect our approach to be compatible for applications to other diseases and types of perturbations, as we have shown already with our genetic dependency model.

Finally, the analyses performed in this study represent a foundation for the development of further, enhanced deep learning models to predict drug response and genetic dependency in pursuit of identifying anti-cancer treatments targeting specific cancer gene expression signatures. While beyond the scope of this study, a fusion model combining the many different modalities and modeling strategies examined in this work is one avenue of further study warranting future exploration. In addition, as more data becomes available, we expect to see performance levels for future versions of our models improve, as we have already seen with the increased performance of our SSeq2.0 MLP and CNN models relative to SSeq1.0 models.

### Author Contributions

B.A.O.: Conceptualization, methodology, software, formal analysis, investigation, implementation, visualization, writing – original draft, and writing – review & editing.

J.H.: Methodology, software, formal analysis, investigation, writing – original draft, and writing – review & editing.

V.S.: Conceptualization, methodology, visualization, and writing – review & editing. M.J.M.: Investigation, generation of prospective validation data

K.L.B.: Methodology, supervision, and resources.

S.C.S.: Conceptualization, methodology, supervision, resources, and writing – review & editing.

## Acknowledgements

This work funded by the National Institutes of Health by grants U54HL127624 (NHLBI, LINCS / BD2K), P30CA240139 (NCI, CCSG), R01LM013391 (NLM, Data Curation), and the State of Florida Biomedical Research Program, Bankhead Coley research and research infrastructure grants 9BC13 and 23B16.

## METHODS

### i. Data Aggregation and Pre-Processing

Publicly available data were collected, standardized, and aggregated from multiple independent sources. Cancer cell line drug sensitivity and genetic dependency datasets were sourced from PharmacoDB drug sensitivity experiments^13^ and Project Achilles CRISPR knockout screens, respectively. PharmacoDB monotherapy drug response data and metadata were accessed and downloaded from the PharmacoDB portal and via the PharmacoGx package for *R*^29^. Project Achilles CRISPR genetic dependency screens were accessed and downloaded from the DepMap 23Q4 Public release^30^.

L1000 Phase 3 level-4 mRNA perturbation data for small molecule perturbagens and CRISPR genetic perturbagens were acquired from the LINCS Data Portal (LDP)^12,31,32^. CCLE baseline gene expression profiles for cancer cell line models were accessed and downloaded from the DepMap portal^14–16^. The 20Q4 release of the DepMap CCLE RNAseq TPM expression data was used to assemble the input data utilized in the initial drug sensitivity models. The 23Q4 release of the DepMap CCLE RNAseq TPM expression data was used to assemble the input data for the Genetic Dependency model, as well as for training the finalized *Gen2* SSeq2.0 models.

Each PharmacoDB monotherapy drug sensitivity experiment consisted of a dose-response curve recorded for one human cell line treated with one small molecule compound. The DSS3, a normalized drug sensitivity score, can be derived from the measured AUC of a dose-response inhibition curve. It is calculated by normalizing the DSS2, which is calculated from the DSS1. The DSS1 is a normalized representation of the dose-response AUC. The DSS3 is a robust measure of drug efficacy and potency, and a higher DSS3 corresponds to a higher drug efficacy and greater potency^28^. The methods and code for calculating the DSS3 for PharmacoDB data can be found on the PharmacoGx Github repository (https://github.com/bhklab/PharmacoGx/). Equations 1-4 can be used to calculate the DSS3.

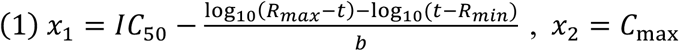

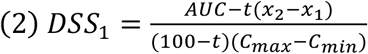

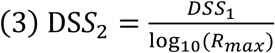

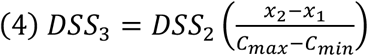

**Equations 1-4.** *C* represents the concentration, *C_max_* represents the maximum dose concentration tested, and *C_min_* represents the minimum concentration tested; (*x_2_–x_1_*) represents the selected concentration range; R_max_ represents the maximal response, or the top asymptote of the dose-response curve; *t* represents the specified minimum activity level, and *b* represents the slope of the curve. *t* was set to 10% as the default value for each experiment.

PharmacoDB dose-response viability normalized, recomputed-AAC and recomputed-IC50 values were used to determine binary categorization of drug response in *SSeq2.0* Gen2 drug sensitivity models. A recomputed-AAC value of 0.275, corresponding to the top 15% of data points, was used as a cut-off to determine sensitivity, with an AAC below 27.5% indicating resistance. Recomputed-IC50 values were incorporated alongside recomputed-AAC values to determine a sensitivity cutoff, with a recomputed-AAC above 27.5% required for an experiment to be labeled as ‘sensitive’.

### Matching Compounds and Cell Lines across Sources

Compounds and cell lines from PharmacoDB datasets were first mapped to LINCS CMap-L1000 compounds and DepMap CCLE cell lines to ensure compatibility across sources. Metadata sourced from PubChem, LINCS Data Portal, and ChEMBL were used to assist in matching small molecule identifiers for compounds present in both PharmacoDB and CMap datasets, while metadata sourced from Cellosaurus were used to assist in matching and standardizing cell line model identifiers across sources for cell lines present in both PharmacoDB and DepMap CCLE. Compound identifiers were mapped across sources using compound metadata from each respective source by cross-referencing database entries contained in ChEMBL, PubChem, and LDP to ensure matching accuracy. Compound metadata and information used for mapping include chemical and structural information, SMILEs, synonyms, and database accession identifiers. Cell lines were mapped across sources by cross-referencing metadata from each source to Cellosaurus entries to ensure data standardization and integrity^33^.

#### Data Filtering Steps

L1000 Phase 3 (December 2021 release) level-4 normalized small molecule perturbation gene expression signature data were used to generate transcription consensus signatures (TCS) for small molecules using methodology adapted from and described in our previous work^22^. Compound data were filtered to retain compound perturbations measured at 24 hours for at least two different dose concentrations and in at least two different cell lines. The methods described in ^22^ were applied to the December 2021 release of level-4 LINCS CMap-L1000 small molecule perturbation signatures for the 978 landmark genes measured at the 24-hour timepoint to aggregate by dose and by cell-line, to generate TCS representative of biological perturbation activity across 230 L1000-measured cell lines. TCS for 7,398 LINCS CMap-L1000 compounds were generated and used as model input for training, development, evaluation, and generating predictions. These 7,398 LINCS CMap-L1000 compounds were later screened *in silico* by using the generated TCS as input in the finalized DNN model to generate predicted activity against 1,287 CCLE cell lines, using the respective compound TCS and CCLE signatures as input, as shown in Figure 1.

L1000 TCS and CCLE expression datasets were each separately scaled to a proportional single unit range of (0, 1) using the *scales* package for R^34^. CCLE RNAseq TPM data were obtained from the DepMap portal, scaled, and filtered to retain only expression values for genes in common with the 978 L1000 landmark genes. This filtering step led to 969 overlapping genes retained for each of our machine learning input datasets. CCLE mass spectrometry protein quantification data were also obtained from the DepMap portal^27^, and the same steps that were followed for the RNAseq data were followed for the protein quantification DNN model.

Monotherapy PharmacoDB experiments composed of one cell line treated with one compound were downloaded via PharmacoGx for *R* and filtered to retain experiments consisting of one CCLE cell line treated with one LINCS CMap-L1000 compound, with a DSS3 value as the measured response variable. PharmacoDB datasets used in this study included CCLE, CTRPv2, GDSC1000, gCSI, GRAY, FIMM, and UHNBreast.

PharmacoDB2.0 datasets used included the same datasets in addition to PRISM and NCI60. Data from each of the datasets were aggregated and filtered to remove duplicate cell line-compound experiments. For instances of duplicate compound-cell line pairs, the experiment with the highest DSS3 or recomputed-AAC value was retained, while the remaining experiments were excluded.

A PharmacoDB-CCLE-CMap L1000 machine learning input dataset was synthesized by aggregating filtered, overlapping, and scaled data collected across three independent sources: PharmacoDB monotherapy drug sensitivity screening datasets, DepMap CCLE baseline gene expression profiles for cancer cell line models, and LINCS CMap-L1000 compound perturbation signatures. The final aggregated data consisted of a combination of transcriptional data for the retained 969 landmark genes for each compound and for each cell line, alongside monotherapy measures of drug sensitivity, in the form of DSS3 values, for each compound-cell line pair tested. The aggregated PharmacoDB-CCLE-L1000 dataset was randomly partitioned into three separate sets for training, validation, and evaluation. Thirty percent of the filtered PharmacoDB-CCLE-CMap L1000 drug sensitivity data was set aside as a fully separate test set to later evaluate the prediction performance of the fully trained model. The 60% training set was used for initial model training, while the 10% validation set was used to assess and improve model performance during training, after each epoch.

A cutoff for determining sensitivity was initially set for DSS3 values greater than 0.04, which corresponded to values less than about 5.0 micromolar (μM) for IC50 and to values greater than 27.5 for recomputed-AAC. Cohen’s kappa, κ, used to assess agreement for binarized IC50 and recomputed-AAC based on cutoff, can be calculated using Equations 5-7.

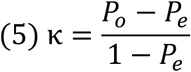

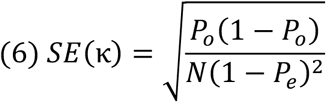

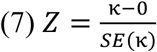

**Equations 5-7**. Cohen’s Kappa, κ, can be calculated using equation 5. Equations 6 and 7 can be used to determine the Z-score and level of significance of the agreement between observations. Where *P_o_* is the observed agreement, *P_e_* is the expected agreement, and *N* is the total number of observations. Each observation corresponds to one cell line-compound viability experiment.

### ii. CRISPR Genetic Dependency Data Integration and Pre-Processing

The methodology employed to generate small molecule TCS was adapted for the CRISPR perturbation signatures for 978 landmark genes measured at the 96-hour timepoint, resulting in the generation of 13,390 CRISPR TCS^35^. Additionally, CCLE cell line RNA-sequencing gene expression profiles (23Q4) and gene dependency data from Project Achilles, encompassing gene dependency probability estimates for all CRISPR knockout screens in the integrated gene effect, were acquired from the DepMap Portal (https://depmap.org/portal/)^30,36^. A low gene dependency score indicates a reduced probability of the cell-line being dependent on the gene, while a high dependency score implies a greater likelihood of reliance^35,37^.

To ensure data integrity, CCLE cell lines were mapped to Achilles cell lines based on metadata standardized and available through the DepMap Portal. LINCS CRISPR perturbation signatures and CCLE RNA-seq TPM profiles were filtered, revealing 969 overlapping genes in common from the 978 L1000 landmark genes. A similar filtering process was applied to genetic dependency data to retain unique samples of cell line and perturbagen combinations. The input dataset for the machine learning model comprised the unique combination of L1000 CRISPR TCS, CCLE cell lines, and binary scores based on gene dependency values.

A gene dependency cutoff of 0.5 was applied, distinguishing the top 10% of scores as positive for genetic dependency, with the remaining 90% of scores categorized as negative for genetic dependency. The input dataset was randomly split into separate sets corresponding to 60:10:30 proportions, allocating 60% for training, 10% for post-training validation, and 30% for testing to evaluate the final model.

### iii. Training and developing deep neural network models

#### PharmacoDB drug sensitivity

Our goal was to develop a model to predict the likelihood of a given cell line exhibiting sensitivity to a given compound, based on transcriptional data for a subset of 969 genes representative of each entity, returning a rank-ordered list of compounds that can be prioritized for further investigation. Each of the two sets of transcriptional signatures was passed into one of two respective dense-layer convolutional neural subnetworks. Following a series of dense layers and a linear activation function in the final layer of each subnetwork, the output of each of the two subnetworks were merged and subsequently connected to another series of dense layers (Figure 3). We employed binary encoding and classification of PharmacoDB normalized drug sensitivity scores (DSS3), with a cutoff of 0.04, to train the model to evaluate and predict potential sensitivity of cell lines to LINCS CMap-L1000 small molecules based on the respective gene expression signatures of each. After training the binary classification model, the test set was used to evaluate the performance and accuracy of the prediction estimates generated by the model.

To analyze prediction estimates from the DNN model, a cutoff of 50% probability of a ‘1’, indicating sensitivity, (0.5) was used. Prediction estimates greater than or equal to 0.5 were categorized as a ‘1’ (sensitive), and prediction estimates less than 0.5 were categorized as ‘0’ (resistant). Using these cutoffs to evaluate the model’s performance in predicting outcomes for our test set, our primary pan-cancer model showed 90.5% accuracy (Table S1).

Given a drug sensitivity experiment, *x*, with a drug sensitivity score, *S*, for a cell line, *c*, treated with a drug or compound, *d*, an experiment can be summarized as

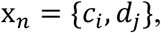

and the sensitivity score can be estimated as a function of the interaction of the gene expression, *G*, of a gene, *n*, in response to perturbation by a drug, *d*, or as part of the cancer profile of a cell line, *c*:

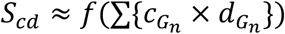

Integrated drug sensitivity data from PharmacoDB, aligned with corresponding gene expression signatures for both compounds and cell lines, were joined to form a labeled dataset that was then randomly split into three separate sets to be used for (1) training (60%), (2) validation (10%), and (3) testing and evaluation (30%). Prior to model input, data were standardized and scaled to a single unit range from 0 to 1 using the rescale function of the *scales* package for *R*^34^.

Various model architectures and training methods were tested to arrive at our finalized model architecture. The finalized model consists of a MLP deep neural network model with two parallel subnetworks containing a series of dense layers, followed by a concatenation layer, and another series of dense layers (Figure 3). Supervised learning was employed to train each binary classification model.

A basic, multivariate logistic regression model was trained and evaluated as a benchmark against which to evaluate our more complex DNN models. A logistic regression model using the drug response dataset used in this study can be summarized using Equation 8.

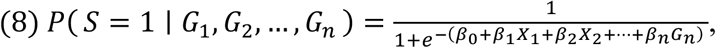

**Equation 8**. Logistic regression can be used to model the drug sensitivity data used in this study to compare performance to more complex models. *P(S=1)* represents the probability of a ‘sensitive’ drug response outcome, *G_n_* represents the gene expression, *G*, of a gene, *n*, as the predictor variables, and *ß_0:n_* represent the model coefficients.

Keras and Tensorflow, via the Keras package for *R*, were used to construct and develop DNN architectures and machine learning models. For each MLP model, we constructed a model with two input MLP dense subnetworks utilizing ReLU activation, with linear activation in the final layer of each subnetwork, followed by a merge layer preceding a final series of dense layers, ending in a single-node layer with sigmoidal activation for binary categorization to (1) classify cell line sensitivity response to a compound, for our drug sensitivity prediction model, or (2) classify gene dependency based on response to genetic perturbation by CRISPR reagent. Binary cross-entropy was employed as the loss function for all models. The Adam optimizer was used for model optimization for each MLP model and each CNN model, along with concurrent validation using the separate 10%-allocation validation set for all models. The predicted probability of the positive class, *ŷ*, can be estimated for the described DNN architectures as

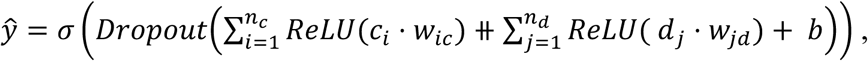

where σ represents the sigmoid activation function applied in the final output layer, *n* represents the number of input nodes in a given dense layer, *w* represents the weights of a given layer, and *b* represents the bias. (⧺ symbolizes concatenation of the two input feature subnetworks for *c* and *d*.)

MLP drug sensitivity models were trained for 25 epochs, while CNN models were trained for 20 epochs. A batch size of 5,000 was employed for training all MLP drug sensitivity models, while a batch size of 2,000 was used to train CNN drug sensitivity models. A batch size of 50,000 was used to train the genetic dependency model. Model performance was assessed after training for each model using a separate 30%-allocation test set, which was set aside prior to training.

### iv. Evaluation of deep neural network models

Multiple evaluation criteria and metrics were used to evaluate the performance of each model. The ROC is a measure of the true positive rate (TPR) as a function of the false positive rate (FPR). The TPR, also known as recall or sensitivity, is the number of correctly predicted sensitive experiments (positive classes) divided by the total number of actual sensitive experiments, and the FPR is the number of incorrectly predicted sensitive experiments (false positive classes) divided by the number of actual resistant experiments:

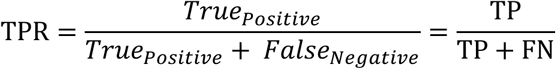

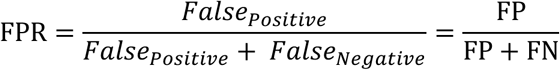

where TP is the number of true positives, FN is the number of false negatives, and FP is the number of false positives.

The ROC curve represents the diagnostic ability of a binary classifier system as its discrimination threshold is varied and is summarized in a single value by measuring the area under the curve (AUROC). The yardstick package for *R* was used to assess ROC and calculate the AUROC of all SSeq models^38^. The yardstick package was also used to measure the precision-recall curve and calculate the AUPR.

Precision, also known as the positive predictive value, is measured as

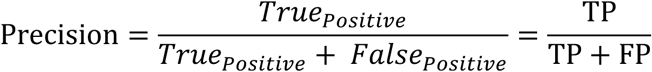

Specificity, or the true negative rate (TNR), and overall accuracy were also measured. Specificity measures the proportion of actual negative classes that are correctly identified by the model, and can be calculated as

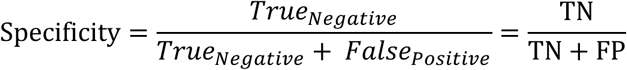

Accuracy measures the proportion of correct predictions out of the total number of predictions made and can be calculated as

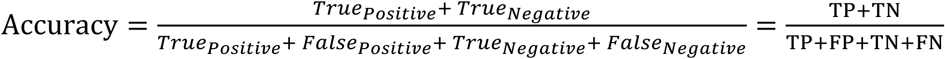

Model tuning and hyperparameters were optimized and validated using Keras Tuner.

### v. Cross-Validation and Stratification

Cross-validation methods were employed to ensure robustness and accuracy of prediction capacity for our finalized model. First, Monte Carlo cross-validation and 10-fold (*k*-fold) cross-validation methods were employed to assess how the finalized model architecture and attributes performed for various subsets of sample data (Tables S3, S4, and S5). Next, tissue-based cross-domain validation was employed using a leave-one-group-out approach to assess prediction accuracy of our methods and model architecture for a tissue type that the model had not been trained on (Extended Data Fig. 1; Table S6). Compound structural data were similarly utilized to facilitate a compound-based cross-domain validation analysis with compounds grouped by FCFP4-20 and ECFP4-10 fingerprint clusters (Extended Data Fig. 1 and Extended Data Fig. 2).

#### ECFP4 Fingerprint Generation and Model Development

L1000 small molecule perturbagen IDs were mapped to canonical smiles in the compound metadata obtained from the LINCS CMap-L1000 data source. After filtering out missing values, ECFP4 descriptors with a bit length of 1024 were computed for the remaining small molecules using RDkit toolkits in Python as described in one of our previous studies^26^. The aggregated L1000 TCS-ECFP4-CCLE-PharmacoDB dataset comprised 421 unique small molecule compounds and 983 unique cell lines, resulting in 286,737 datapoints with DSS3 labels. Using L1000 TCS, CCLE, and ECFP4 as inputs, we created an MLP model containing an additional, third input feature subnetwork, parallel to the existing two feature subnetworks in the SSeq MLP model, and with identical in construction to the existing subnetworks, except for a 1024-unit input layer to accommodate the dimensions of the ECFP4 fingerprint data. We also created an ECFP4-CCLE MLP model by replacing the L1000 TCS input features with our ECFP4 input features. This was accomplished by altering the 969-unit input layer to a 1024-unit input layer and feeding the ECFP4 data into the new model alongside CCLE gene expression signatures for training and evaluation.

### vi. External, Prospective Validation in Prostate Cancer

#### Cell line RNA-sequencing and *in vitro* drug screens

RNAseq reads were obtained for eight prostate human cell lines: seven PC cell line models (*22Rv1*, *C4-2B, DU145*, *LNCaP Clone FGC*, *NCI-H660, PC3*, and *VCaP*) and one non-tumorigenic prostate epithelial cell line (*RWPE-1).* Cell lines were originally obtained from American Type Culture Collection (ATCC) and were authenticated for contamination by LabCorp prior to culturing.

LNCaP Clone FGC (LNCaP), 22Rv1, DU145 and PC3 cells were cultured in RPMI (Corning, 15-040-CV) supplemented with 100 mg/mL penicillin/streptomycin (ThermoFisher, 15140122), 2 mM L-glutamine (ThermoFisher, 25030024), and 10% fetal bovine serum (FBS) (Atlanta Biologicals). C4-2B cells were cultured in DMEM (Corning, 10-013-CV) supplemented as described above. VCaP cells were cultured in DMEM GlutaMax (Gibco, 10569-044) and supplemented with 1% anti-anti (ThermoFisher, 15240062) and 10% FBS. NCI-H660 were cultured in RPMI-1640 high glucose (ATCC, 30-2001) supplemented with 1% penicillin/streptomycin, 2mM L-glutamine (additional), 1% Insulin Transferrin-Selenium (Gibco, 41400-045), 10nM Hydrocortisone (Sigma-Aldrich, H0135), 10nM β-estradiol (Sigma-Aldrich, E2257), and 5% FBS. RWPE-1 cells were cultured in Keratinocyte-SFM (ThermoFisher, 17005042) supplemented with 1% penicillin/streptomycin, 25mg Bovine Pituitary Extract (Gibco, 13028-014), and 2.5 μg human recombinant EGF (Gibco, 10450-013). For drug treatment assays, LNCaP and VCaP cells were cultured in 2% FBS and C4-2B, 22Rv1, PC3, DU145 and NCI-H660 cells were cultured in 2% charcoal-stripped serum (CSS). For RWPE-1 and VCaP cells, drugs were screened in the same media as their maintenance growth conditions. Cell cultures were maintained at 37°C in a humidified atmosphere of 5% CO_2_.

Total RNA from the eight prostate cell lines models was harvested using TRIzol (Thermo Fisher) and purified using Direct-zol RNA MiniPrep Plus reagent (Zymo Research), according to the manufacturer’s protocol. In total 24 samples were sequenced (including 3 replicates for each of the eight cell lines) and more than 30M reads were produced for each sample. FASTQ files were examined and processed for quality control using FastQC^39^. Cutadapt was used to trim raw reads to remove Illumina adapters and low-quality reads^40^. A quality score of 20 was used as a cutoff to trim reads, and sequences shorter than 20bp were discarded. Filtered sequences were aligned to the hg19 human reference genome using STAR aligner. Reads were quantified into Transcripts per Million (TPM) and normalized using the DESeq2 package for R^41^. Transcripts were mapped to HGNC gene symbols based on hg19 human reference genome annotation metadata.

Normalized TPM data were scaled to a range from 0 to 1 using the *scales* package for R^34^ and filtered for the 969 landmark genes used to train pan-cancer ML models to generate gene expression signatures reflective of PC cell lines. Filtered prostate cell line RNAseq TPM were input into the SSeq2.0 MLP drug sensitivity model (in place of CCLE RNAseq TPM) alongside L1000 TCS for 7,398 compounds to generate drug sensitivity predictions.

Using PC RNAseq TPM as input, drug sensitivity predictions were generated for PC cell lines to 7,398 LINCS CMap-L1000 compounds selected using criteria described prior using the SSeq2.0 MLP model. Out of the 7,398 compounds included in the analysis, 208 compounds were predicted to be active against one or more PC cell lines. From the prioritized list of compounds predicted to be active against PC cell lines, 26 commercially available small molecule compounds were selected for further experimental investigation based on availability and FDA approval or investigational drug status. Eleven of 26 drugs were selected based on existing FDA approval status, one compound (rotenone) has undergone preclinical investigation for use in cancer^42^, while the rest selected were clinical investigational drugs. Additionally, 13 of the 26 drugs selected have undergone at least the first phase of clinical trials but have not reached FDA approval.

Twenty-six compounds with available L1000 TCS were selected for *in vitro* screening on the eight prostate cell lines, and drug treatment assays were carried out using the methods described in our previous study^43^. Briefly, RWPE-1, 22Rv1, PC3, DU145 (2,500 cells/well), LNCaP (1,000 cells/well), C4-2B (5,000 cells/well), NCI-H660 (30,000 cells/well) and VCaP (50,000 cells/well) were seeded in 96 well plates (white microclear, Greiner) and incubated overnight at 37 C. The following day, cells were treated with varying drug concentrations (0 μM, 0.37 μM, 1.11 μM, 3.33 μM, 10 μM, and 30 μM) using the OT-2 liquid handling robot (Opentron). Cells were imaged every 4 h using the Incucyte ZoomSystem for 72 h and cell viability was measured using Cell-Titer Glo end-point assay. Luminescence was read using Glo-Max plate reader (Promega). IC50 values were calculated using the GR calculator online tool^44^.

Predictions were evaluated using binarized IC50 values from PC experimental sensitivity data, using a cutoff of IC50 < 5.0 µM indicating sensitivity. SSeq1.0 was evaluated on the same dataset as SSeq2.0 for performance comparison.

### Reproducibility

A seed was set before all computational work and analyses performed in this study. For reproducibility, a seed of 1234 was set for computational work, as described, except where stated otherwise. For model repetitions, seeds 111, 222, 123, 1234, and 12345 were used to produce five replicates per model unless indicated otherwise. Code and data used in this study are available as described in Code Availability and Data Availability, respectively. Keras models were developed in Python 3.7 and in *R* using TensorFlow backend (version 2.4.0-GPU) in Python 3.7, via the Keras *R* package (version 2.11.1).

### Lead Contact

Further information and requests for resources and reagents should be directed to and will be fulfilled by the lead contact and corresponding author, Stephan Schurer.

### Materials Availability

This study did not generate new unique reagents.

### Code Availability

Code for the deep learning models and Shiny application is available via the Schurer Lab SensitivitySeq project GitHub page (https://github.com/schurerlab/sensitivityseq)..

### Data Availability

All training data used are available through public online sources. LINCS CMap-L1000 data can be accessed via the LINCS Data Portal (https://lincsportal.ccs.miami.edu/datasets/view/LDS-1611). Level-4 CMap L1000 Phase 3 2021 small molecule data are available from LINCS Data Portal (https://identifiers.org/lincs.data/LDS-1613) where they were retrieved, and are also available from CLUE (https://clue.io/data/CMap2020#LINCS2020).

CCLE RNAseq TPM profiles can be accessed from the Dependency Map Project Portal (DepMap, Broad (2020): DepMap 20Q4 Public (https://doi.org/10.6084/m9.figshare.13681534.v1), and DepMap, Broad (2023): DepMap 23Q4 Public (https://doi.org/10.25452/figshare.plus.24667905.v2).

PharmacoDB sensitivity data are available from PharmacoDB (https://pharmacodb.pmgenomics.ca/) and the PharmacoGx package for *R*^29^. Standardized datasets used in this study have been deposited to Zenodo and NCBI Sequence Read Archive (SRA). PC model input data are available via Zenodo: https://doi.org/10.5281/zenodo.13329140. PC RNAseq FASTQ files have been deposited to NCBI SRA (BioProject accession: PRJNA1149944).

Prospective human prostate cell line sensitivity data used in this study are provided along with other model data in Supplemental Data and deposited data. *SensitivitySeq* predictions for L1000 compounds and CCLE cell lines are available for query via the *SensitivitySeq* application (SensitivitySeq.com). All other data supporting the findings of this study are available via the *SensitivitySeq* Github repository (https://github.com/schurerlab/sensitivityseq/) and upon request from the corresponding author.

### Supplemental information

Document S1. Extended Data Fig. 1–3 and Tables S1–S11.

Table S12. External Prostate Cancer Predictions/Validation Table, related to Table S11.

