## Supplementary material for "Pan-Cancer Drug Sensitivity Prediction from Gene Expression using Deep Learning": Document S1. Supplementary Data

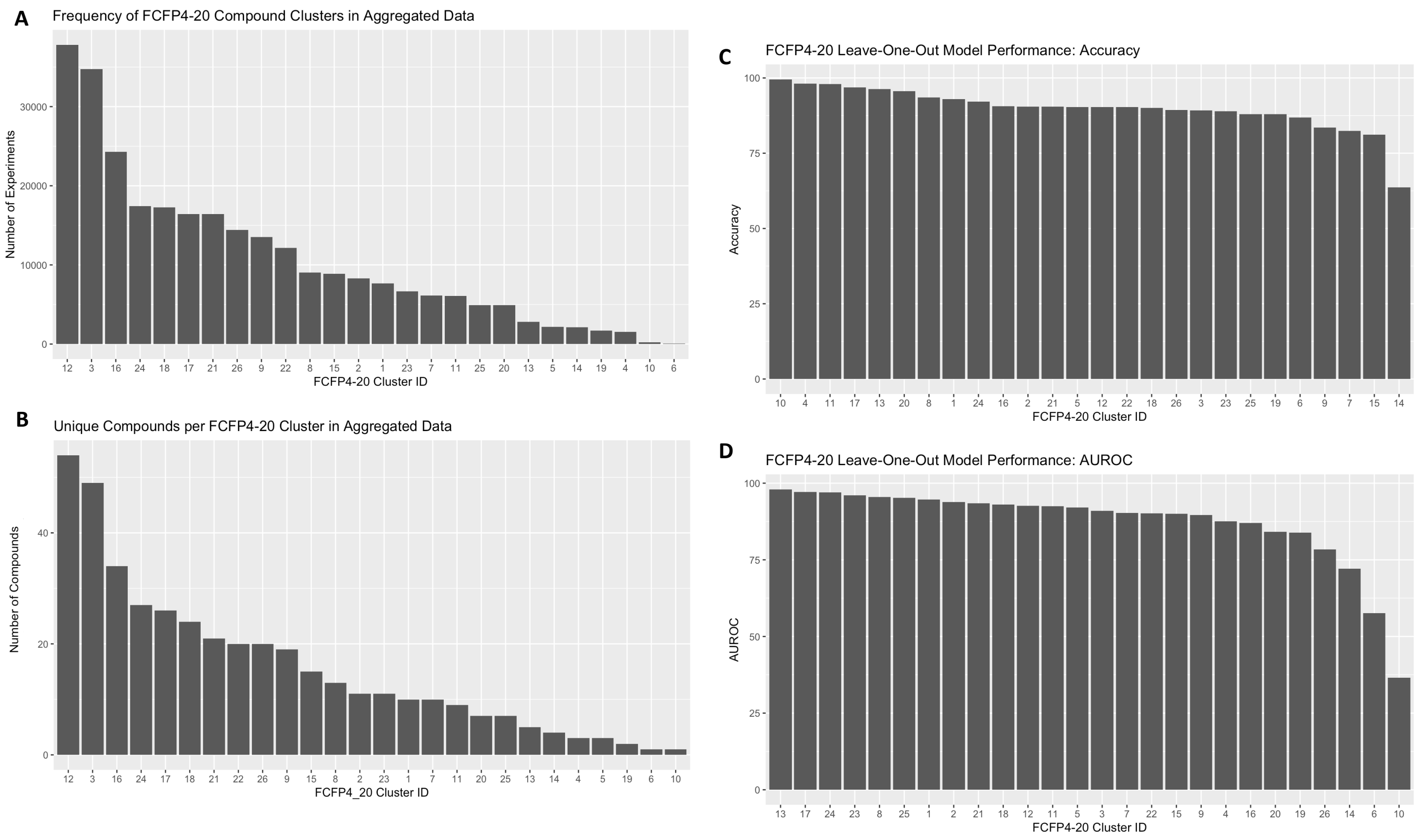


**Extended Data Fig. 1. Small molecule compound groups clustered by FCFP4-20 fingerprints distribution and compound cross-domain validation model performance results.** (A) Distribution of compound groups clustered by FCFP4-20 fingerprints in aggregated model development dataset. (B) Representation of FCFP4-20 fingerprint compound clusters in aggregated model development dataset. (C) Compound FCFP4-20 fingerprint cluster cross-validation leave-one out model performance accuracy by fingerprint cluster group. (D) Compound FCFP4-20 fingerprint cluster cross-validation leave-one out model performance AUROC by fingerprint cluster group.

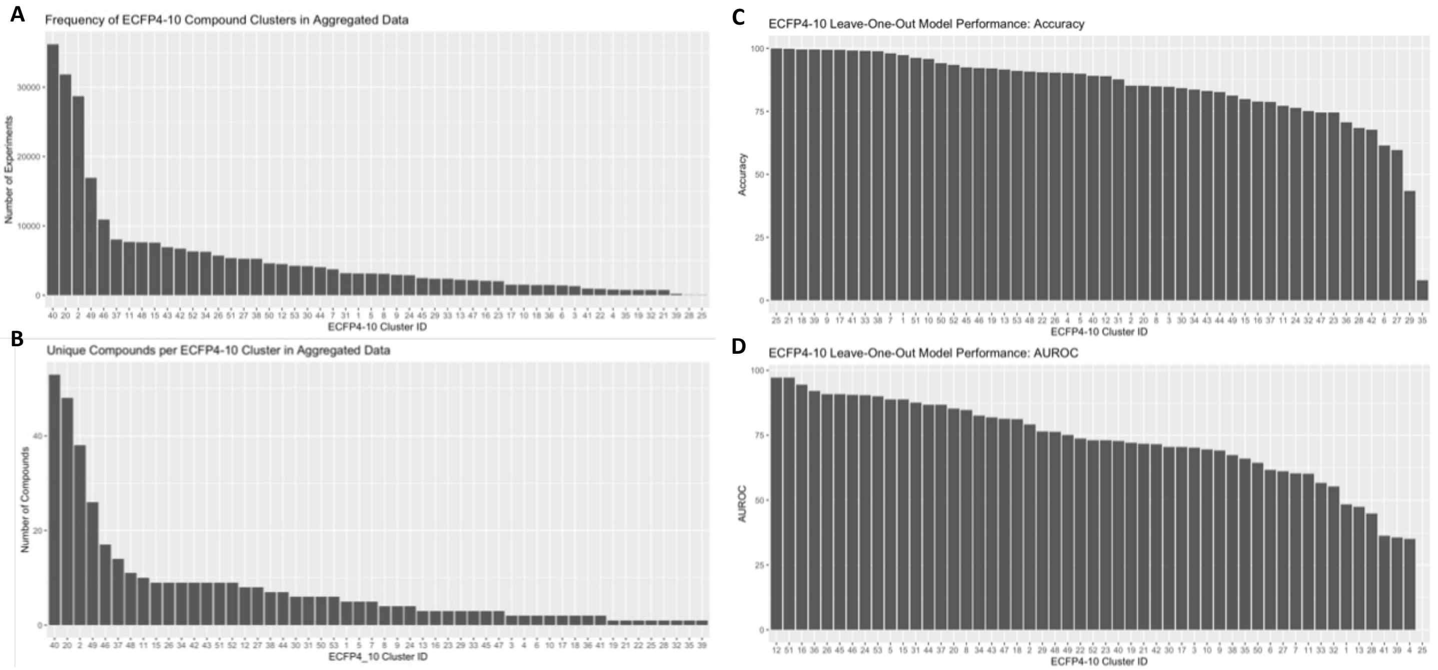


**Extended Data Fig. 2. Small molecule compound groups clustered by ECFP4-10 fingerprints distribution and compound cross-domain validation model performance results.** (A) Distribution of compound groups clustered by ECFP4-10 fingerprints in aggregated model development dataset. (B) Representation of ECFP4-10 fingerprint compound clusters in aggregated model development dataset. (C) Compound ECFP4-10 fingerprint cluster cross-validation leave-one out model performance accuracy by fingerprint cluster group. (D) Compound ECFP4-10 fingerprint cluster cross-validation leave-one out model performance AUROC by fingerprint cluster group.

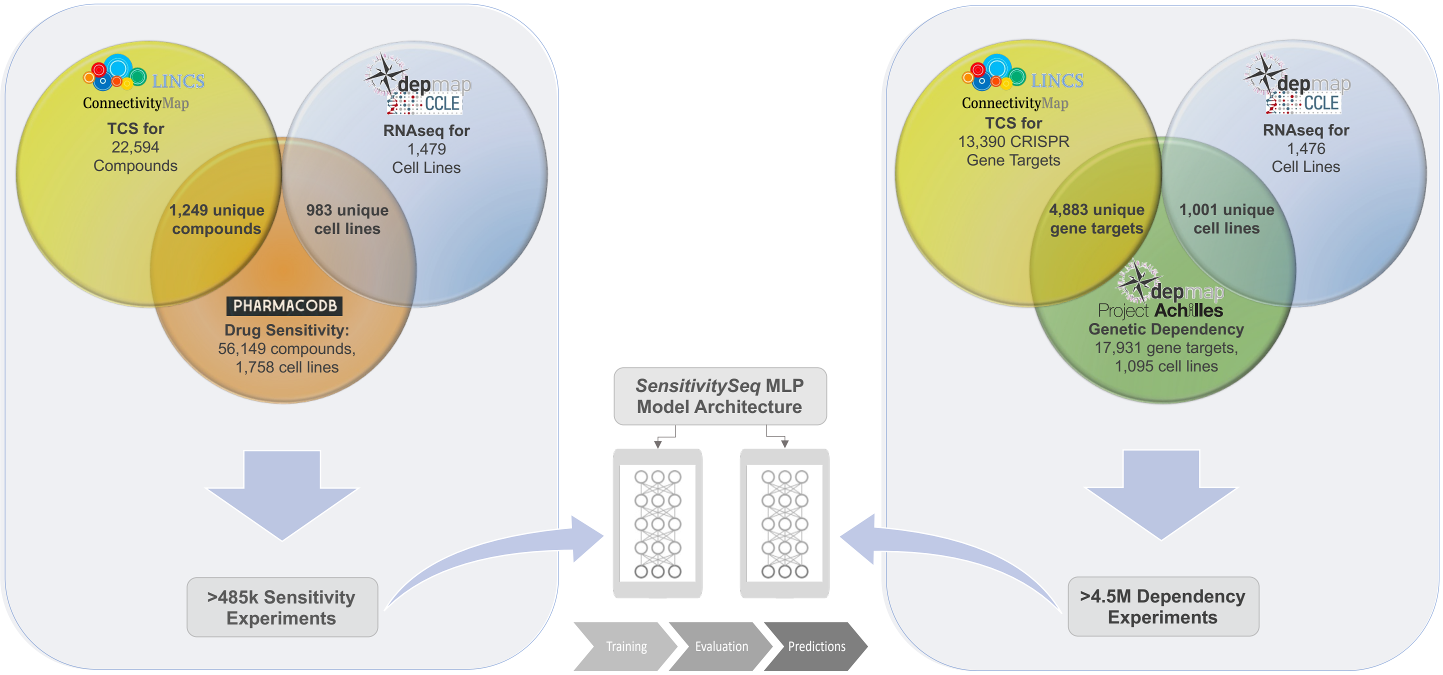


**Extended Data Fig. 3. Abstract illustration summarizing aggregated datasets used for training *SensitivitySeq2.0* models.** Large, aggregated datasets were used to train drug sensitivity and genetic dependency SSeq2.0 models. The *Gen2* drug sensitivity training set consisted of nearly 500,000 unique experiments across 1,249 small molecule compounds and 983 cancer cell lines, while the genetic dependency dataset spanned 4,883 CRISPR gene targets and 1,001 cancer cell lines.

| Model | Loss | Accuracy | AUROC | AUPR | Precision | Recall | Specificity |
|---|---|---|---|---|---|---|---|
| **Drug Sensitivity MLP (pre-scaling)** | 0.2367 | 89.87% | 92.40% | 73.08% | 70.60% | 58.80% | 95.50% |
| ***SSeq1.0* DS MLP** | 0.2264 | 90.54% | 93.07% | 76.17% | 73.46% | 62.84% | 95.74% |
| **Full CCLE Transcriptome DS MLP** | 0.2410 | 89.89% | 92.04% | 74.10% | 73.90% | 55.74% | 96.30% |
| **Randomized Labels** | 0.4375 | 84.20% | 48.96% | 18.53% | 0.00% | undefined | 100.0% |

**Table S1. Summary of evaluation and performance for models tested during initial model development.** The first model represents the performance for our initial pan-cancer Drug Sensitivity (DS) MLP model, trained and evaluated prior to scaling input datasets. The SSeq1.0 DS MLP model in row 2 reflects the performance for the primary, pan-cancer L1000-CCLE-PharmacoDB MLP model trained with scaled input data from LINCS CMap-L1000 compound TCS and CCLE gene expression signatures. The Full CCLE Transcriptome DS MLP represents the performance metrics for a model developed using the full, transcriptome-wide set of genes present in the CCLE RNAseq data as cancer cell line input features, in contrast to the 969-gene subset of the CCLE data filtered for only the L1000 landmark genes that was used for the other models. The Randomized Labels model corresponds to the SSeq1.0 MLP drug sensitivity model architecture and input features trained with ‘scrambled’ or randomly reordered outcome labels (while maintaining the imbalanced class ratio).

| Model | Loss | Accuracy | AUROC | AUPR | Precision | Recall | Specificity |
|---|---|---|---|---|---|---|---|
| ***SSeq1.0* DS MLP** | 0.2264 | 90.54% | 93.07% | 76.17% | 73.46% | 62.84% | 95.74% |
| **DS MLP, CW** | 0.3412 | 84.91% | 93.03% | 75.70% | 51.33% | 87.64% | 84.40% |
| **DS 1D-CNN** | 0.2420 | 89.97% | 92.01% | 73.15% | 72.39% | 59.10% | 95.77% |
| **DS 2D-CNN** | 0.2935 | 89.10% | 91.22% | 72.84% | 63.85% | 71.57% | 92.39% |
| ***SSeq1.0* DS 2D-CNN, CW** | 0.2691 | 88.84% | 93.44% | 76.94% | 61.08% | 80.93% | 90.32% |
| **Logistic Regression** | 0.4215 | 84.20% | 66.29% | 28.87% | 0.00% | undefined | 100.0% |

**Table S2. Summary of DNN Model Performance for Various Model Structures, related to Figure 6.** Initial drug sensitivity (DS) model predictions were evaluated using several measures of performance as criteria. The first model listed represents the performance for our initial primary pan-cancer drug sensitivity model, SSeq1.0, as a point of comparison. An equivalent architecture to SSeq1.0 with the addition of 5:1 class weight optimization (CW) was evaluated, but this balancing strategy did not lead to better overall performance than SSeq1.0 without CW. The 2D- and 1D-CNN models represent additional early models that were trained, validated, and evaluated using the same datasets as our initial Drug Sensitivity MLP model. The 1D-CNN was not retained further due to suboptimal recall performance. The 2D-CNN architecture was evaluated with the addition of CW, leading to the retention of the CW 2D-CNN as a finalized SSeq1.0 model. In the final row, performance is shown for a basic logistic regression model trained and evaluated on the same data as prior models in the table.

| Model Architecture | Evaluation | Rep | Training Allocation (%) | Testing Allocation (%) | Seed | Loss | Accuracy | AUROC | Precision | Recall | Specificity |
|---|---|---|---|---|---|---|---|---|---|---|---|
| SSeq MLP | 70:30 Sample Split Set 1 | 1 | 70 | 30 | 123 | 0.2387 | 89.87% | 92.31% | 71.06% | 58.26% | 95.66% |
| SSeq MLP | 70:30 Sample Split Set 2 | 2 | 70 | 30 | 1234 | 0.2367 | 89.87% | 92.39% | 72.55% | 55.44% | 96.17% |
| SSeq MLP | 70:30 Sample Split Set 3 | 3 | 70 | 30 | 12345 | 0.2461 | 89.59% | 92.12% | 67.29% | 63.47% | 94.36% |
| SSeq MLP | 70:30 Sample Split Set 4 | 4 | 70 | 30 | 123456 | 0.2399 | 89.94% | 92.26% | 72.10% | 56.90% | 95.98% |
| SSeq MLP | 70:30 Sample Split Set 5 | 5 | 70 | 30 | 654321 | 0.2443 | 89.82% | 92.15% | 70.09% | 60.06% | 95.29% |
| SSeq MLP | 70:30 Sample Split Sets 1-5 Mean | 1-5 | 70 | 30 | N/A | 0.2411 | 89.82% | 92.25% | 70.62% | 58.83% | 95.49% |
| SSeq MLP | 70:30 Sample Split Sets 1-5 Std Dev | 1-5 | 70 | 30 | N/A | 0.0039 | 0.1352% | 0.1135% | 2.093% | 3.104% | 0.7129% |

**Table S3. Monte Carlo repeated random sub-sampling cross-validation with 70% of aggregated data allocated to training.** Model performance results for Monte Carlo repeated random sub-sampling cross-validation models with 70% of aggregated data allocated to training are summarized.

| Model Architecture | Evaluation | Rep | Training Allocation (%) | Testing Allocation (%) | Seed | Loss | Accuracy | AUROC | Precision | Recall | Specificity |
|---|---|---|---|---|---|---|---|---|---|---|---|
| SSeq MLP | 1:99 Sample Split Set 1 | 1 | 1 | 99 | 123 | 0.3203 | 87.92% | 86.21% | 64.73% | 48.36% | 95.17% |
| SSeq MLP | 1:99 Sample Split Set 2 | 2 | 1 | 99 | 1234 | 0.3349 | 87.56% | 86.58% | 64.35% | 44.07% | 95.53% |
| SSeq MLP | 1:99 Sample Split Set 3 | 3 | 1 | 99 | 12345 | 0.3074 | 87.59% | 86.54% | 62.13% | 50.75% | 94.34% |
| SSeq MLP | 1:99 Sample Split Set 4 | 4 | 1 | 99 | 123456 | 0.3034 | 87.30% | 86.39% | 61.82% | 46.98% | 94.69% |
| SSeq MLP | 1:99 Sample Split Set 5 | 5 | 1 | 99 | 654321 | 0.3024 | 87.81% | 86.10% | 65.44% | 44.85% | 95.67% |
| SSeq MLP | 1:99 Sample Split Sets 1-5 Mean | 1-5 | 1 | 99 | N/A | 0.3137 | 87.64% | 86.37% | 63.69% | 47.00% | 95.08% |
| SSeq MLP | 1:99 Sample Split Sets 1-5 Std Dev | 1-5 | 1 | 99 | N/A | 0.01385 | 0.2406% | 0.2086% | 1.620% | 2.698% | 0.5620% |

**Table S4. Monte Carlo repeated random sub-sampling cross-validation with 1% of aggregated data allocated to training.** Model performance results for Monte Carlo repeated random sub-sampling cross-validation models with 1% of aggregated data allocated to training are summarized.

| Model Architecture | Evaluation | Rep | Training Allocation (%) | Testing Allocation (%) | Seed | Loss | Accuracy | AUROC | Precision | Recall | Specificity |
|---|---|---|---|---|---|---|---|---|---|---|---|
| SSeq MLP | Sample Set 1 | 1 | 10 | 90 | 123 | 0.2695 | 88.98% | 90.66% | 68.95% | 52.32% | 95.69% |
| SSeq MLP | Sample Set 2 | 2 | 10 | 90 | 123 | 0.2740 | 88.53% | 90.40% | 76.58% | 37.31% | 97.91% |
| SSeq MLP | Sample Set 3 | 3 | 10 | 90 | 123 | 0.2811 | 88.38% | 90.43% | 62.65% | 61.68% | 93.27% |
| SSeq MLP | Sample Set 4 | 4 | 10 | 90 | 123 | 0.2677 | 88.76% | 90.67% | 68.49% | 50.39% | 95.77% |
| SSeq MLP | Sample Set 5 | 5 | 10 | 90 | 123 | 0.2700 | 89.01% | 90.87% | 68.20% | 54.49% | 95.34% |
| SSeq MLP | Sample Set 6 | 6 | 10 | 90 | 123 | 0.2704 | 88.62% | 90.87% | 63.60% | 61.68% | 93.55% |
| SSeq MLP | Sample Set 7 | 7 | 10 | 90 | 123 | 0.2755 | 88.91% | 90.70% | 68.37% | 52.91% | 95.51% |
| SSeq MLP | Sample Set 8 | 8 | 10 | 90 | 123 | 0.2716 | 88.93% | 90.71% | 67.55% | 54.84% | 95.17% |
| SSeq MLP | Sample Set 9 | 9 | 10 | 90 | 123 | 0.2665 | 88.78% | 90.73% | 67.46% | 53.22% | 95.30% |
| SSeq MLP | Sample Set 10 | 10 | 10 | 90 | 123 | 0.2690 | 88.98% | 90.81% | 68.27% | 53.82% | 95.42% |
| SSeq MLP | Sample Sets 1-10 Mean | 1-10 | 10 | 90 | 123 | 0.2715 | 88.79% | 90.69% | 68.01% | 53.27% | 95.29% |
| SSeq MLP | Samples Set 1-10 Std Dev | 1-10 | 10 | 90 | 123 | 0.0043 | 0.2167% | 0.1615% | 3.703% | 6.748% | 1.268% |

**Table S5. 10-fold cross-validation results.** Model performance results are shown for 10-fold *k-fold* cross-validation models. Ten unique slices of 10% of the total aggregated data were allocated as training sets with the remaining 90% of data used for evaluation.

| CCLE Tissue Type | Sample Size | Loss | Accuracy | AUROC | AUPR | Precision | Recall | Specificity |
|---|---|---|---|---|---|---|---|---|
| ADRENAL CORTEX | 147 | 0.2834 | 89.80% | 90.09% | 67.28% | 63.33% | 82.61% | 91.13% |
| AUTONOMIC GANGLIA | 5792 | 0.2862 | 87.62% | 91.24% | 73.57% | 72.43% | 57.08% | 94.85% |
| BILIARY TRACT | 2493 | 0.2245 | 89.93% | 91.59% | 62% | 61.57% | 48.53% | 95.75% |
| BREAST | 17150 | 0.2764 | 88.14% | 89.83% | 75.67% | 68.29% | 51.07% | 95.37% |
| BONE | 6097 | 0.2688 | 88.81% | 90.95% | 66.60% | 74.06% | 58.75% | 95.45% |
| CENTRAL NERVOUS SYSTEM | 17580 | 0.2212 | 90.52% | 92.40% | 70.34% | 79.59% | 36.97% | 98.57% |
| CERVIX | 2110 | 0.3032 | 86.30% | 89.11% | 66.73% | 62.23% | 60.42% | 91.97% |
| ENDOMETRIUM | 9087 | 0.2169 | 90.93% | 93.81% | 74.57% | 62.70% | 74.19% | 93.43% |
| FIBROBLAST | 2194 | 0.2069 | 90.70% | 87.55% | 43.50% | 40.44% | 43.79% | 94.62% |
| HAEMATOPOIETIC AND LYMPHOID TISSUE | 52836 | 0.3149 | 86.71% | 90.66% | 78.79% | 80.20% | 56.71% | 95.77% |
| KIDNEY | 7504 | 0.2295 | 90.37% | 90.29% | 57.29% | 56.30% | 58.53% | 94.33% |
| LARGE INTESTINE | 17088 | 0.2583 | 88.99% | 91.13% | 66.94% | 60.77% | 66.02% | 92.84% |
| LIVER | 7608 | 0.2439 | 89.56% | 90.65% | 62.46% | 61.93% | 57.31% | 94.55% |
| LUNG | 54175 | 0.2267 | 90.61% | 92.17% | 71.60% | 69.99% | 59.54% | 95.76% |
| OESOPHAGUS | 9935 | 0.2124 | 91.27% | 93.66% | 75.37% | 73.83% | 60.13% | 96.45% |
| OVARY | 14738 | 0.2135 | 90.98% | 93.39% | 72.93% | 69.27% | 62.84% | 95.51% |
| PANCREAS | 13444 | 0.1977 | 91.37% | 93.61% | 70.48% | 66.92% | 58.38% | 95.97% |
| PLACENTA | 312 | 0.3137 | 85.26% | 85.60% | 35.41% | 39.53% | 45.95% | 90.55% |
| PLEURA | 2875 | 0.2083 | 91.55% | 92.19% | 65.66% | 65.37% | 56.06% | 96.15% |
| PROSTATE | 1791 | 0.2817 | 88.44% | 89.97% | 69.41% | 73.30% | 52.26% | 96.02% |
| SALIVARY GLAND | 672 | 0.2222 | 89.58% | 93.49% | 74.75% | 57.41% | 72.09% | 92.15% |
| SKIN | 17169 | 0.2211 | 90.89% | 91.80% | 67.86% | 68.80% | 54.53% | 96.31% |
| SMALL INTESTINE | 183 | 0.4873 | 78.14% | 88.17% | 76.26% | 81.82% | 44.26% | 95.08% |
| SOFT TISSUE | 8701 | 0.2227 | 91.09% | 93.51% | 79.30% | 77.65% | 64.74% | 96.31% |
| STOMACH | 11224 | 0.2272 | 90.32% | 92.56% | 71.98% | 76.11% | 48.40% | 97.43% |
| THYROID | 4607 | 0.2198 | 91.19% | 93.39% | 76.23% | 81.86% | 51.76% | 98.01% |
| UPPER AERODIGESTIVE TRACT | 10916 | 0.2154 | 91.21% | 93.48% | 74.60% | 75.84% | 58.35% | 96.83% |
| URINARY TRACT | 8451 | 0.2025 | 91.91% | 93.70% | 74.84% | 75.06% | 59.75% | 96.91% |
| Mean | 10959.96 | 0.2502 | 89.36% | 91.43% | 68.66% | 67.74% | 57.18% | 95.15% |
| Std Dev | 13275.33 | 0.0583 | 2.778% | 2.099% | 9.836% | 10.72% | 9.615% | 1.975% |

**Table S6. Cell line cross-domain validation model performance by CCLE Tissue Type, related to Figure 4.**

| Ligand Type | Loss | Accuracy | AUROC | Precision | Recall | Specificity | Unique Compounds | Test Set Size |
|---|---|---|---|---|---|---|---|---|
| Activator | 0.3135 | 89.34% | 93.29% | 84.06% | 93.43% | 86.14% | 7 | 8939 |
| Agonist | 0.1561 | 95.74% | 87.79% | 80.88% | 56.47% | 98.92% | 18 | 13108 |
| Antagonist | 0.2523 | 88.84% | 42.36% | 1.63% | 3.04% | 92.41% | 17 | 10738 |
| Blocker[a] | 0.0126 | 100% | 100.0% | 100.0% | 100.0% | 100.0% | 1 | 36 |
| Enhancer | 0.1284 | 96.77% | 93.93% | 81.58% | 83.78% | 98.08% | 3 | 804 |
| Inhibitor | 0.5633 | 81.50% | 75.73% | 48.77% | 59.95% | 86.22% | 355 | 325799 |
| Ligand—other/unknown | 0.3006 | 90.50% | 71.27% | 0% | undefined | 100.0% | 1 | 926 |
| Modulator | 0.1004 | 98.82% | 35.00% | 0% | undefined | 100.0% | 1 | 929 |
| Stimulant | 0.4078 | 79.98% | 87.00% | 69.21% | 99.37% | 64.34% | 3 | 2123 |
| Stabilizing Agent | 0.7027 | 57.56% | 42.84% | 58.46% | 97.39% | 0.00% | 2 | 714 |
| Mean | 0.2938 | 87.91% | 72.92% | 52.46% | 74.18% | 82.61% | 40.80 | 36411.60 |
| Std Dev | 0.2155 | 12.65% | 24.26% | 38.49% | 33.55% | 31.08% | 110.59 | 101795.74 |

[a]no positive classes present in test set

**Table S7. Cross-domain validation by compound ligand category.**

| Model | Loss | Accuracy | AUROC | AUPR | Precision | Recall | Specificity |
|---|---|---|---|---|---|---|---|
| **Balanced by Exclusion (*SSeq1.0* MLP)** | 0.3734 | 84.14% | 92.59% | 73.94% | 49.91% | 87.71% | 83.48% |
| **Balanced by Fusion (*SSeq1.0* MLP)** | 0.3313 | 85.88% | 93.34% | 76.42% | 53.24% | 87.37% | 85.59% |
| ***SSeq1.0* DS MLP, CW** | 0.3412 | 84.91% | 93.03% | 75.70% | 51.33% | 87.64% | 84.40% |

**Table S8. Model performance after applying various balancing strategies, related to Figure 6.** Model performance results are shown following application of three balancing strategies tested. An exclusion-based balancing strategy (row 1), in which training set experiments with negative (resistant) outcomes were randomly excluded until reaching approximately equal, 1:1 proportions of experiments with resistant outcomes to those with sensitive outcomes. In row 2, a fusion-based balancing strategy was applied by randomly splitting the resistant-outcome training set experiments into five groups, with each subset approximately proportional to the sensitive-outcome training set experiments subset. Subsequently, each resistant-outcome subset was separately combined with the sensitive-outcome subset to form five smaller, balanced training data subsets. Training subsets were used consecutively to train a fusion MLP model in a series of five training steps, for 5 epochs per subset to equal 25 total training epochs. In row 3, a class-weights, hyperparameter-based strategy was used to train each model with 5:1 class weights set for sensitive:resistant classes. Setting class weights to 5:1 leads to each sensitive-outcome experiment exerting the same level of influence on model weights as five resistant-outcome experiments during training. All models were evaluated using the same separate, unaltered, and untransformed test set.

| Model | Loss | Accuracy | AUROC | AUPR | Precision | Recall | Specificity |
|---|---|---|---|---|---|---|---|
| ***SSeq1.0* CCLE-only Input MLP** | 0.4304 | 84.20% | 60.00% | 22.18% | 0.00% | undefined | 100.0% |
| ***SSeq1.0* L1000-only Input MLP** | 0.2540 | 89.32% | 91.38% | 70.21% | 71.24% | 54.35% | 95.88% |
| **CCLE Protein Quantification MLP** | 0.2229 | 90.65% | 93.09% | 74.50% | 70.61% | 65.40% | 95.15% |
| ***SSeq2.0* Genetic Dependency MLP** | 0.1670 | 92.86% | 98.44% | 90.49% | 55.28% | 95.44% | 92.61% |

**Table S9. Comparison of DNN Models for Various Input Features and Datasets, related to Figure 6.**

| Model | Loss | Accuracy | AUROC | AUPR | Precision | Recall | Specificity |
|---|---|---|---|---|---|---|---|
| **TCS, TPM Dual-Subnetwork MLP** | 0.2532 | 89.49% | 91.36% | 73.10% | 75.55% | 53.38% | 96.60% |
| **ECFP4, TCS, TPM 3-Subnetwork MLP** | 0.2330 | 90.10% | 93.25% | 76.74% | 78.46% | 54.95% | 97.03% |
| **ECFP4, TPM Dual-Subnetwork MLP** | 0.2379 | 89.57% | 93.13% | 76.16% | 82.20% | 46.73% | 98.01% |

**Table S10. Comparison of performance for drug sensitivity models trained on an ECFP4-annotated subset of the initial aggregated dataset.**

| Model | Accuracy | AUROC | AUPR | Precision | Recall | Specificity | True Positives | True Negatives | False Positives | False Negatives |
|---|---|---|---|---|---|---|---|---|---|---|
| **PC Validation, *SSeq1.0* MLP** | 59.87% | 70.72% | 88.30% | 83.70% | 57.42% | 67.09% | 267 | 106 | 52 | 198 |
| **PC Validation, *SSeq2.0* MLP** | 67.42% | 65.87% | 85.83% | 74.17% | 86.45% | 11.39% | 402 | 18 | 140 | 63 |

**Table S11. Prospective, external PC validation of *SensitivitySeq*.** SSeq drug sensitivity MLP models were validated in prostate cancer (PC) cell lines. SSeq2.0 was validated using a prospective, single-tissue experimental validation. SSeq1.0 was also externally validated using the same dataset for comparison. RNAseq TPM for eight cell lines with three biological replicates each were used as input to generate predictions for each cell line for each L1000 TCS compound available. Following *in vitro* validation experiments, a 5.0 μM IC50 cutoff was applied to determine actual classes and evaluate predicted classes.
