## Supplementary material for "Pan-Cancer Drug Sensitivity Prediction from Gene Expression using Deep Learning": Table S12

| Cell line | Compound | IC50 (actual) | Actual score | SSeq1.0 prediction estimate | SSeq1.0 predicted score | SSeq2.0 prediction estimate | SSeq2.0 predicted score |
|---|---|---|---|---|---|---|---|
| 22Rv1 | afatinib | 1.88 | 1 | 0.142782360 | 0 | 0.538164198 | 1 |
| C4-2B | afatinib | 5.45 | 0 | 0.149451077 | 0 | 0.690935493 | 1 |
| DU145 | afatinib | 2.25 | 1 | 0.149314344 | 0 | 0.542400181 | 1 |
| LNCaP | afatinib | 1.22 | 1 | 0.071935117 | 0 | 0.689906955 | 1 |
| NCI-H660 | afatinib | 7.82 | 0 | 0.021622747 | 0 | 0.538164198 | 1 |
| PC3 | afatinib | 3.95 | 1 | 0.206208199 | 0 | 0.561330438 | 1 |
| RWPE-1 | afatinib | 0.09 | 1 | 0.384666979 | 0 | 0.538164198 | 1 |
| VCaP | afatinib | 23.6 | 0 | 0.049034894 | 0 | 0.539431930 | 1 |
| 22Rv1 | alisertib | 30 | 0 | 0.412455201 | 0 | 0.558874667 | 1 |
| C4-2B | alisertib | 30 | 0 | 0.562531710 | 1 | 0.711332798 | 1 |
| DU145 | alisertib | 13.45 | 0 | 0.217999965 | 0 | 0.558874667 | 1 |
| LNCaP | alisertib | 20.59 | 0 | 0.355319440 | 0 | 0.705398202 | 1 |
| NCI-H660 | alisertib | 30 | 0 | 0.586519897 | 1 | 0.558874667 | 1 |
| PC3 | alisertib | 16.94 | 0 | 0.107419848 | 0 | 0.560064554 | 1 |
| RWPE-1 | alisertib | 10.06 | 0 | 0.268422067 | 0 | 0.558874667 | 1 |
| VCaP | alisertib | 30 | 0 | 0.297763586 | 0 | 0.560448468 | 1 |
| 22Rv1 | alvocidib | 0.06 | 1 | 0.917779446 | 1 | 0.897899508 | 1 |
| C4-2B | alvocidib | 0 | 1 | 0.936987877 | 1 | 0.957896471 | 1 |
| DU145 | alvocidib | 0 | 1 | 0.878314674 | 1 | 0.901111364 | 1 |
| LNCaP | alvocidib | 0 | 1 | 0.894602180 | 1 | 0.957792997 | 1 |
| NCI-H660 | alvocidib | 30 | 0 | 0.822813869 | 1 | 0.897899508 | 1 |
| PC3 | alvocidib | 0 | 1 | 0.807090402 | 1 | 0.915017545 | 1 |
| RWPE-1 | alvocidib | 0 | 1 | 0.908599854 | 1 | 0.897899508 | 1 |
| VCaP | alvocidib | 0 | 1 | 0.826730430 | 1 | 0.900076807 | 1 |
| 22Rv1 | anisomycin | 0 | 1 | 0.535566986 | 1 | 0.780103922 | 1 |
| C4-2B | anisomycin | 0 | 1 | 0.609309793 | 1 | 0.899583340 | 1 |
| DU145 | anisomycin | 0.01 | 1 | 0.393346429 | 0 | 0.783417702 | 1 |
| LNCaP | anisomycin | 0.01 | 1 | 0.485879332 | 0 | 0.899318457 | 1 |
| NCI-H660 | anisomycin | 30 | 0 | 0.427598000 | 0 | 0.780103922 | 1 |
| PC3 | anisomycin | 0.08 | 1 | 0.283856273 | 0 | 0.805271149 | 1 |
| RWPE-1 | anisomycin | 0.01 | 1 | 0.433127880 | 0 | 0.780103922 | 1 |
| VCaP | anisomycin | 5.12 | 0 | 0.429514557 | 0 | 0.780978084 | 1 |
| 22Rv1 | BI-2536 | 0 | 1 | 0.943608165 | 1 | 0.874229670 | 1 |
| C4-2B | BI-2536 | 0 | 1 | 0.958806992 | 1 | 0.947653890 | 1 |
| DU145 | BI-2536 | 0 | 1 | 0.910637140 | 1 | 0.878232718 | 1 |
| LNCaP | BI-2536 | 0 | 1 | 0.925457656 | 1 | 0.947536826 | 1 |
| NCI-H660 | BI-2536 | 4.45 | 1 | 0.883519888 | 1 | 0.874229670 | 1 |
| PC3 | BI-2536 | 0 | 1 | 0.811250925 | 1 | 0.894525409 | 1 |
| RWPE-1 | BI-2536 | 0 | 1 | 0.935109138 | 1 | 0.874229670 | 1 |
| VCaP | BI-2536 | 0 | 1 | 0.876823068 | 1 | 0.876939058 | 1 |
| 22Rv1 | cabazitaxel | 30 | 0 | 0.725094736 | 1 | 0.955084205 | 1 |
| C4-2B | cabazitaxel | 0 | 1 | 0.769216478 | 1 | 0.982285380 | 1 |
| DU145 | cabazitaxel | 0 | 1 | 0.679298043 | 1 | 0.956856966 | 1 |
| LNCaP | cabazitaxel | 0 | 1 | 0.684690356 | 1 | 0.982240796 | 1 |
| NCI-H660 | cabazitaxel | 30 | 0 | 0.514086246 | 1 | 0.955084205 | 1 |
| PC3 | cabazitaxel | 0 | 1 | 0.614686549 | 1 | 0.963878632 | 1 |
| RWPE-1 | cabazitaxel | 0 | 1 | 0.710921526 | 1 | 0.955084205 | 1 |
| VCaP | cabazitaxel | 0 | 1 | 0.597662449 | 1 | 0.956288099 | 1 |
| 22Rv1 | cladribine | 0.31 | 1 | 0.518917680 | 1 | 0.752118945 | 1 |
| C4-2B | cladribine | 0.12 | 1 | 0.578958452 | 1 | 0.878757596 | 1 |
| DU145 | cladribine | 1.85 | 1 | 0.367065161 | 0 | 0.755814433 | 1 |
| LNCaP | cladribine | 0.14 | 1 | 0.410308838 | 0 | 0.878445268 | 1 |
| NCI-H660 | cladribine | 180 | 0 | 0.276999652 | 0 | 0.752118945 | 1 |
| PC3 | cladribine | 22.39 | 0 | 0.266851485 | 0 | 0.752966881 | 1 |
| RWPE-1 | cladribine | 0.28 | 1 | 0.438352138 | 0 | 0.752118945 | 1 |
| VCaP | cladribine | 30 | 0 | 0.345904648 | 0 | 0.753245115 | 1 |
| 22Rv1 | danusertib | 30 | 0 | 0.07794854 | 0 | 0.767408252 | 1 |
| C4-2B | danusertib | 0 | 1 | 0.091372281 | 0 | 0.890387833 | 1 |
| DU145 | danusertib | 1.22 | 1 | 0.048509479 | 0 | 0.770642757 | 1 |
| LNCaP | danusertib | 0 | 1 | 0.071571082 | 0 | 0.890101612 | 1 |
| NCI-H660 | danusertib | 30 | 0 | 0.087356687 | 0 | 0.767408252 | 1 |
| PC3 | danusertib | 45.37 | 0 | 0.029868245 | 0 | 0.768267870 | 1 |
| RWPE-1 | danusertib | 0.83 | 1 | 0.053665698 | 0 | 0.767408252 | 1 |
| VCaP | danusertib | 30 | 0 | 0.077460647 | 0 | 0.768509865 | 1 |
| 22Rv1 | dasatinib | 24.63 | 0 | 0.672664464 | 1 | 0.587255418 | 1 |
| C4-2B | dasatinib | 11.85 | 0 | 0.680742025 | 1 | 0.742329836 | 1 |
| DU145 | dasatinib | 0.21 | 1 | 0.683224559 | 1 | 0.591964960 | 1 |
| LNCaP | dasatinib | 14.86 | 0 | 0.583333731 | 1 | 0.738205552 | 1 |
| NCI-H660 | dasatinib | 30 | 0 | 0.148270190 | 0 | 0.587255418 | 1 |
| PC3 | dasatinib | 22.74 | 0 | 0.652873993 | 1 | 0.588304222 | 1 |
| RWPE-1 | dasatinib | 0.48 | 1 | 0.708958447 | 1 | 0.587255418 | 1 |
| VCaP | dasatinib | 30 | 0 | 0.293738723 | 0 | 0.588645995 | 1 |
| 22Rv1 | delanzomib | 3.31 | 1 | 0.515109599 | 1 | 0.944406748 | 1 |
| C4-2B | delanzomib | 0 | 1 | 0.597142160 | 1 | 0.978011966 | 1 |
| DU145 | delanzomib | 2.98 | 1 | 0.368071824 | 0 | 0.946456015 | 1 |
| LNCaP | delanzomib | 0.25 | 1 | 0.461424947 | 0 | 0.977956831 | 1 |
| NCI-H660 | delanzomib | 0.04 | 1 | 0.396050096 | 0 | 0.944406748 | 1 |
| PC3 | delanzomib | 5.48 | 0 | 0.260596097 | 0 | 0.944864273 | 1 |
| RWPE-1 | delanzomib | 0 | 1 | 0.425655454 | 0 | 0.944406748 | 1 |
| VCaP | delanzomib | 0 | 1 | 0.403796315 | 0 | 0.945749938 | 1 |
| 22Rv1 | docetaxel | 0 | 1 | 0.743937314 | 1 | 0.805332005 | 1 |
| C4-2B | docetaxel | 0 | 1 | 0.759613454 | 1 | 0.915802717 | 1 |
| DU145 | docetaxel | 0 | 1 | 0.734765053 | 1 | 0.810435236 | 1 |
| LNCaP | docetaxel | 0 | 1 | 0.704077244 | 1 | 0.915620685 | 1 |
| NCI-H660 | docetaxel | 30 | 0 | 0.272650301 | 0 | 0.805332005 | 1 |
| PC3 | docetaxel | 0 | 1 | 0.699551582 | 1 | 0.832469940 | 1 |
| RWPE-1 | docetaxel | 0 | 1 | 0.754433155 | 1 | 0.805332005 | 1 |
| VCaP | docetaxel | 0 | 1 | 0.562275946 | 1 | 0.808715940 | 1 |
| 22Rv1 | doxorubicin | 1.76 | 1 | 0.917484283 | 1 | 0.933674097 | 1 |
| C4-2B | doxorubicin | 2.32 | 1 | 0.937280893 | 1 | 0.973536968 | 1 |
| DU145 | doxorubicin | 2.67 | 1 | 0.873442113 | 1 | 0.935979366 | 1 |
| LNCaP | doxorubicin | 0.7 | 1 | 0.891810477 | 1 | 0.973470867 | 1 |
| NCI-H660 | doxorubicin | 3.52 | 1 | 0.825995564 | 1 | 0.933674097 | 1 |
| PC3 | doxorubicin | 2.61 | 1 | 0.796601772 | 1 | 0.945927560 | 1 |
| RWPE-1 | doxorubicin | 0.25 | 1 | 0.905400634 | 1 | 0.933674097 | 1 |
| VCaP | doxorubicin | 1.83 | 1 | 0.822135806 | 1 | 0.935229301 | 1 |
| 22Rv1 | foretinib | 0 | 1 | 0.520031989 | 1 | 0.626131713 | 1 |
| C4-2B | foretinib | 3.94 | 1 | 0.614345908 | 1 | 0.776917577 | 1 |
| DU145 | foretinib | 1.15 | 1 | 0.352992982 | 0 | 0.630936027 | 1 |
| LNCaP | foretinib | 0 | 1 | 0.458470374 | 0 | 0.776169538 | 1 |
| NCI-H660 | foretinib | 12 | 0 | 0.467473716 | 0 | 0.626131713 | 1 |
| PC3 | foretinib | 0.94 | 1 | 0.245098054 | 0 | 0.649018824 | 1 |
| RWPE-1 | foretinib | 1.8 | 1 | 0.408249021 | 0 | 0.626131713 | 1 |
| VCaP | foretinib | 0.168 | 1 | 0.400226921 | 0 | 0.629600048 | 1 |
| 22Rv1 | irinotecan | 1.09 | 1 | 0.521790862 | 1 | 0.499655545 | 0 |
| C4-2B | irinotecan | 7.7 | 0 | 0.601659656 | 1 | 0.657574058 | 1 |
| DU145 | irinotecan | 2.09 | 1 | 0.377734035 | 0 | 0.506911993 | 1 |
| LNCaP | irinotecan | 2.08 | 1 | 0.466879964 | 0 | 0.655854404 | 1 |
| NCI-H660 | irinotecan | 30 | 0 | 0.404456347 | 0 | 0.499655545 | 0 |
| PC3 | irinotecan | 73.7 | 0 | 0.269218087 | 0 | 0.499655545 | 0 |
| RWPE-1 | irinotecan | 2.94 | 1 | 0.431501299 | 0 | 0.499655545 | 0 |
| VCaP | irinotecan | 0.21 | 1 | 0.405478597 | 0 | 0.504889667 | 1 |
| 22Rv1 | lestaurtinib | 1.02 | 1 | 0.631889641 | 1 | 0.720049024 | 1 |
| C4-2B | lestaurtinib | 0 | 1 | 0.664161921 | 1 | 0.857274532 | 1 |
| DU145 | lestaurtinib | 0.66 | 1 | 0.616400003 | 1 | 0.725078881 | 1 |
| LNCaP | lestaurtinib | 0.16 | 1 | 0.607629836 | 1 | 0.856915832 | 1 |
| NCI-H660 | lestaurtinib | 16.95 | 0 | 0.323508173 | 0 | 0.720049024 | 1 |
| PC3 | lestaurtinib | 0.99 | 1 | 0.486676842 | 0 | 0.741457582 | 1 |
| RWPE-1 | lestaurtinib | 0.34 | 1 | 0.621448755 | 1 | 0.720049024 | 1 |
| VCaP | lestaurtinib | 30 | 0 | 0.479613513 | 0 | 0.721600533 | 1 |
| 22Rv1 | mitoxantrone | 0.15 | 1 | 0.999408245 | 1 | 0.775885582 | 1 |
| C4-2B | mitoxantrone | 0.3 | 1 | 0.999579608 | 1 | 0.896502733 | 1 |
| DU145 | mitoxantrone | 0 | 1 | 0.998866916 | 1 | 0.778867543 | 1 |
| LNCaP | mitoxantrone | 0.08 | 1 | 0.999170661 | 1 | 0.896283746 | 1 |
| NCI-H660 | mitoxantrone | 1.77 | 1 | 0.998643041 | 1 | 0.775885582 | 1 |
| PC3 | mitoxantrone | 0.68 | 1 | 0.997340024 | 1 | 0.799872756 | 1 |
| RWPE-1 | mitoxantrone | 0.22 | 1 | 0.999204516 | 1 | 0.775885582 | 1 |
| VCaP | mitoxantrone | 0.66 | 1 | 0.998522580 | 1 | 0.777858913 | 1 |
| 22Rv1 | MK-1775 | 1.53 | 1 | 0.309861064 | 0 | 0.564128220 | 1 |
| C4-2B | MK-1775 | 0.62 | 1 | 0.371170461 | 0 | 0.717569947 | 1 |
| DU145 | MK-1775 | 0.93 | 1 | 0.216849953 | 0 | 0.567659974 | 1 |
| LNCaP | MK-1775 | 0.66 | 1 | 0.256115049 | 0 | 0.716593683 | 1 |
| NCI-H660 | MK-1775 | 1335.2 | 0 | 0.305694252 | 0 | 0.564128220 | 1 |
| PC3 | MK-1775 | 1.11 | 1 | 0.143258214 | 0 | 0.588397324 | 1 |
| RWPE-1 | MK-1775 | 20.97 | 0 | 0.251462638 | 0 | 0.564128220 | 1 |
| VCaP | MK-1775 | 30 | 0 | 0.252946675 | 0 | 0.565177321 | 1 |
| 22Rv1 | MLN-0128 | 0 | 1 | 0.680450320 | 1 | 0.469782561 | 0 |
| C4-2B | MLN-0128 | 0 | 1 | 0.715497196 | 1 | 0.639645100 | 1 |
| DU145 | MLN-0128 | 0 | 1 | 0.654029965 | 1 | 0.469782561 | 0 |
| LNCaP | MLN-0128 | 0 | 1 | 0.651003242 | 1 | 0.637790620 | 1 |
| PC3 | MLN-0128 | 0 | 1 | 0.572612405 | 1 | 0.515787244 | 1 |
| RWPE-1 | MLN-0128 | 0 | 1 | 0.668436527 | 1 | 0.469782561 | 0 |
| VCaP | MLN-0128 | 30 | 0 | 0.581978798 | 1 | 0.476449728 | 0 |
| 22Rv1 | panobinostat | 0.04 | 1 | 0.997586012 | 1 | 0.971533060 | 1 |
| C4-2B | panobinostat | 0.03 | 1 | 0.998270631 | 1 | 0.988816619 | 1 |
| DU145 | panobinostat | 0 | 1 | 0.995521784 | 1 | 0.972703636 | 1 |
| LNCaP | panobinostat | 0 | 1 | 0.996653855 | 1 | 0.988788247 | 1 |
| NCI-H660 | panobinostat | 0 | 1 | 0.994546354 | 1 | 0.971533060 | 1 |
| PC3 | panobinostat | 0.01 | 1 | 0.989740372 | 1 | 0.977001488 | 1 |
| RWPE-1 | panobinostat | 0.17 | 1 | 0.996853232 | 1 | 0.971533060 | 1 |
| VCaP | panobinostat | 30 | 0 | 0.994156480 | 1 | 0.971885502 | 1 |
| 22Rv1 | PD-0325901 | 30 | 0 | 0.528346896 | 1 | 0.485723734 | 0 |
| C4-2B | PD-0325901 | 30 | 0 | 0.557128012 | 1 | 0.648250937 | 1 |
| DU145 | PD-0325901 | 5.22 | 0 | 0.554656148 | 1 | 0.500000000 | 1 |
| LNCaP | PD-0325901 | 30 | 0 | 0.344728231 | 0 | 0.641564846 | 1 |
| NCI-H660 | PD-0325901 | 19.7 | 0 | 0.074462116 | 0 | 0.485723734 | 0 |
| PC3 | PD-0325901 | 30 | 0 | 0.610787153 | 1 | 0.487786978 | 0 |
| RWPE-1 | PD-0325901 | 0 | 1 | 0.677488804 | 1 | 0.485723734 | 0 |
| VCaP | PD-0325901 | 30 | 0 | 0.167334914 | 0 | 0.492406249 | 0 |
| 22Rv1 | rotenone | 0 | 1 | 0.530090094 | 1 | 0.611292601 | 1 |
| C4-2B | rotenone | 0 | 1 | 0.621801853 | 1 | 0.763253093 | 1 |
| DU145 | rotenone | 0.244 | 1 | 0.361981481 | 0 | 0.616725922 | 1 |
| LNCaP | rotenone | 0 | 1 | 0.476940334 | 0 | 0.762495756 | 1 |
| NCI-H660 | rotenone | 40.4 | 0 | 0.486687064 | 0 | 0.611292601 | 1 |
| PC3 | rotenone | 0 | 1 | 0.251586825 | 0 | 0.635373592 | 1 |
| RWPE-1 | rotenone | 0.0943 | 1 | 0.418896914 | 0 | 0.611292601 | 1 |
| VCaP | rotenone | 26.3 | 0 | 0.415680200 | 0 | 0.612936735 | 1 |
| 22Rv1 | TAK-733 | 30 | 0 | 0.508037686 | 1 | 0.618616700 | 1 |
| C4-2B | TAK-733 | 30 | 0 | 0.561559737 | 1 | 0.770852149 | 1 |
| DU145 | TAK-733 | 11.9 | 0 | 0.324154258 | 0 | 0.618616700 | 1 |
| LNCaP | TAK-733 | 30 | 0 | 0.288563192 | 0 | 0.766088843 | 1 |
| NCI-H660 | TAK-733 | 30 | 0 | 0.138903528 | 0 | 0.618616700 | 1 |
| PC3 | TAK-733 | 27.08 | 0 | 0.259862334 | 0 | 0.619811952 | 1 |
| RWPE-1 | TAK-733 | 0 | 1 | 0.437957048 | 0 | 0.618616700 | 1 |
| VCaP | TAK-733 | 30 | 0 | 0.219015837 | 0 | 0.620191157 | 1 |
| 22Rv1 | taselisib | 2.05 | 1 | 0.608039081 | 1 | 0.266572446 | 0 |
| C4-2B | taselisib | 0.93 | 1 | 0.624543667 | 1 | 0.527704298 | 1 |
| DU145 | taselisib | 3.81 | 1 | 0.589558959 | 1 | 0.266572446 | 0 |
| LNCaP | taselisib | 0.17 | 1 | 0.508957148 | 1 | 0.525076687 | 1 |
| NCI-H660 | taselisib | 0.62 | 1 | 0.186033905 | 0 | 0.266572446 | 0 |
| PC3 | taselisib | 1.67 | 1 | 0.410802335 | 0 | 0.267707437 | 0 |
| RWPE-1 | taselisib | 0.32 | 1 | 0.604354382 | 1 | 0.266572446 | 0 |
| VCaP | taselisib | 1.02 | 1 | 0.313981295 | 0 | 0.268074363 | 0 |
| 22Rv1 | teniposide | 0.741 | 1 | 0.650498807 | 1 | 0.714690924 | 1 |
| C4-2B | teniposide | 2.97 | 1 | 0.760117710 | 1 | 0.853410304 | 1 |
| DU145 | teniposide | 0 | 1 | 0.401435852 | 0 | 0.719863176 | 1 |
| LNCaP | teniposide | 1.04 | 1 | 0.585485756 | 1 | 0.853043497 | 1 |
| NCI-H660 | teniposide | 0.29 | 1 | 0.678818345 | 1 | 0.714690924 | 1 |
| PC3 | teniposide | 0.601 | 1 | 0.215558469 | 0 | 0.736995697 | 1 |
| RWPE-1 | teniposide | 0.118 | 1 | 0.448394448 | 0 | 0.714690924 | 1 |
| VCaP | teniposide | 4.8 | 1 | 0.506501496 | 1 | 0.718193054 | 1 |
| 22Rv1 | tivantinib | 0.3 | 1 | 0.369169384 | 0 | 0.668236256 | 1 |
| C4-2B | tivantinib | 0.77 | 1 | 0.490735978 | 0 | 0.814820290 | 1 |
| DU145 | tivantinib | 0.45 | 1 | 0.179183096 | 0 | 0.671793818 | 1 |
| LNCaP | tivantinib | 0.34 | 1 | 0.302020311 | 0 | 0.814270616 | 1 |
| NCI-H660 | tivantinib | 30 | 0 | 0.530208409 | 1 | 0.668236256 | 1 |
| PC3 | tivantinib | 30 | 0 | 0.072216481 | 0 | 0.669039249 | 1 |
| RWPE-1 | tivantinib | 0.09 | 1 | 0.218029916 | 0 | 0.668236256 | 1 |
| VCaP | tivantinib | 0.31 | 1 | 0.256141901 | 0 | 0.670608759 | 1 |
| 22Rv1 | vorinostat | 1.66 | 1 | 0.249474913 | 0 | 0.364392698 | 0 |
| C4-2B | vorinostat | 0.85 | 1 | 0.305279911 | 0 | 0.583018184 | 1 |
| DU145 | vorinostat | 0.88 | 1 | 0.161108434 | 0 | 0.364392698 | 0 |
| LNCaP | vorinostat | 0.84 | 1 | 0.211882234 | 0 | 0.581072569 | 1 |
| NCI-H660 | vorinostat | 1.62 | 1 | 0.297014743 | 0 | 0.364392698 | 0 |
| PC3 | vorinostat | 4.86 | 1 | 0.103281438 | 0 | 0.365933567 | 0 |
| RWPE-1 | vorinostat | 1.32 | 1 | 0.181291431 | 0 | 0.364392698 | 0 |
| VCaP | vorinostat | 2.69 | 1 | 0.207778007 | 0 | 0.366428792 | 0 |
